# Population genomics of *Aspergillus sojae* is shaped by the food environment

**DOI:** 10.1101/2023.09.07.556736

**Authors:** Kimberly L. Acevedo, Elizabeth Eaton, Shu Zhao, Katherine Chacon-Vargas, Colin M. McCarthy, Dasol Choi, Jae-Hyuk Yu, John G. Gibbons

## Abstract

Traditional fermented foods often contain specialized microorganisms adapted to the unique food environment. For example, the filamentous mold *Aspergillus oryzae*, used in saké fermentation, has evolved to thrive in starch-rich conditions compared to its wild ancestor, *Aspergillus flavus*. Similarly, *Aspergillus sojae* is used in soybean-based food fermentations (*e.g.* miso and shochu) and is closely related to *Aspergillus parasiticus*. Here, we investigated the impact of long-term *A. sojae* usage in soybean fermentation on population structure, genome variation, and phenotypic traits. We analyzed 12 *A. sojae* and 10 *A. parasiticus* genomes, along with phenotypic characteristics of 15 isolates. Our results revealed that *A. sojae* isolates formed a distinct population separate from *A. parasiticus* and displayed remarkably low levels of genetic diversity, indicative of a recent clonal expansion. Genome comparisons revealed numerous loss-of-function mutations in *A. sojae*, notably in genes responsible for secondary metabolite production, including genes in the aflatoxin encoding gene cluster. Consequently, *A. sojae* lacked aflatoxin production, while it varied among *A. parasiticus* isolates. No other significant differences were observed in growth rates or other measured phenotypic traits between *A. sojae* and *A. parasiticus*. These findings suggest that *A. sojae* may have evolved from a population of *A. parasiticus* and lost the ability to produce some secondary metabolites. To elucidate the phenotypic differences between *A. sojae* and *A. parasiticus*, future work should focus on the influence of wild and food-associated strains on the sensory aspects and microbial community dynamics of fermented soy products.

**Significance Statement:** Like plants and animals, microbes were also domesticated by humans, however relatively little is known about how the process of domestication shapes microbial genomes and traits. We found that isolates of *Aspergillus sojae*, a mold used in the production of miso and soy sauce, makeup a less toxic group that is genetically distinct from its closely related wild ancestor *Aspergillus parasiticus*. Our analyses shed new light on commonalities observed across filamentous molds adapted to the food environment.

## Introduction

Domestication is an evolutionary process in which a population is genetically modified through breeding strategies in an effort to improve traits that are desired by humans (Purugganan & Fuller 2009). For instance, early agriculturalists used selective breeding to select for plants with more food (*i.e.* larger/more seeds and fruits) and livestock that were less aggressive and more fertile (Larson & Fuller 2014). In parallel with plant and animal domestication, humans also unwittingly domesticated microbes (*i.e.* bacteria, yeast and molds) through the continuous propagation of microbial communities in fermented foods (Steensels et al. 2019). This process, referred to as backslopping, resulted in microbial communities that became specialized for their role in improving the longevity, digestibility, and palatability of foods and resulted in genetic differentiation from their ancestral populations (Gibbons & Rinker 2015). For instance, *Lactobacillus bulgaricus* is used as a starter culture in the production of yogurt and has evolved the ability to efficiently metabolize lactose (Sørensen et al. 2016) and casein (Gilbert et al. 1996), the most abundant sugar and protein found in milk, respectively.

Fungi have also played an important historical role in the production of fermented foods and show signatures of specialization to the food environment. For example, *Saccharomyces cerevisiae* shows genetic structure strongly associated with usage (*i.e.* beer, wine, bread etc.) (Legras et al. 2007; Gallone et al. 2016), and lineages have become highly specialized. For example, compared to *S. cerevisiae* wine strains, beer strains can efficiently utilize maltose, a sugar source unique to beer fermentation (Gallone et al. 2016). In addition to yeasts, filamentous fungi have also been domesticated. *Penicillium roqueforti* populations used in the production of blue cheese are differentiated from populations found in food spoilage and natural environments, and two horizontally transferred regions in the blue-cheese populations confer fitness advantages in the dairy environment (Dumas et al. 2020; Cheeseman et al. 2014; Ropars et al. 2015, 2017). Additionally, *Penicillium camemberti*, used in the production of soft cheeses, shows signatures of domestication including population differentiation of cheese populations and phenotypes tailored to the cheese environment (*e.g.* faster growth on cheese in cave conditions, loss of pigmentation, reduced toxin production *etc.*) (Ropars et al. 2020).

The filamentous fungal genus *Aspergillus* additionally includes organisms specialized to the food environment (Gibbons et al. 2012; Sato et al. 2011; Futagami et al. 2011). For instance, *Aspergillus oryzae* has been used in the production of fermented soy based-foods (*e.g.* doenjang and miso) and rice based foods (*e.g.* makgeolli, amazake, and sake) for thousands of years (McGovern et al. 2004; Liu et al. 2019; Machida et al. 2008) and has adapted to the food environment. *A. oyrzae* populations are genetically distinct from their wild progenitor species, *Aspergillus flavus*, and have evolved several adaptations beneficial to the fermentation environment, including increased amylase production to break down starch during rice fermentation, and a reduction or loss of toxin production which may aid in the composition of the microbial community (Gibbons et al. 2012; Watarai et al. 2019; Chacón-Vargas et al. 2021).

*Aspergillus sojae* (AS) is primarily used for soy fermentation because of its high proteolytic activity (Sato et al. 2011; Kim et al. 2019). AS also produces high amounts of leucine aminopeptidase (LAP), which improves flavor development during soybean fermentation (Nampoothiri et al. 2005; Kim et al. 2017). Similar to the *A. oryzae*/*A. flavus* model, it is hypothesized that AS represents a population of *Aspergillus parasiticus* (AP) that is specialized to the food environment (Chang & Hua 2023). AP is an aflatoxin producer and contaminant of stored seeds, grains, and nuts, while AS is a considered as a USDA Generally Regarded As Safe (GRAS) species (Chang et al. 2007). Despite the industrial importance of AS, very little is known about the relationship and phenotypic differences between AS and AP. Here, we analyzed the genomes and characteristics of 22 AS and AP isolates to shed light on their population structure, and genomic and phenotypic differences.

## Methods

### Fungal isolates, culturing, DNA extraction, and Illumina sequencing

AP and AS isolates were obtained from the USDA NRRL ARS culture collection. These include AP NRRL 81, 147, 255, 1032, 1039, 2039, 2040 and 2642 and AS NRRL 70, 299, 1120, 1122, 1123, 1124, 2091, 3351, and 5597. Isolates were cultured on potato dextrose agar (PDA) at 30°C for 48 h. DNA was extracted directly from spores following the protocol described by Zhao *et al*. 2019. The Qubit was used to quantify DNA concentrations for each extraction. PCR-free 150-bp paired-end libraries were constructed and sequenced by Novogene (https://en.novogene.com/) on an Illumina NovaSeq 6000. Raw Illumina whole-genome sequencing data for each strain is available through the BioProject accession number PRJNA911610.

### Illumina sequence quality filtering, read mapping, and variant calling

In addition to the strains sequenced, we also analyzed the publicly available Illumina DNA sequencing data for AP CBS-117618 (NCBI SRA accession number SRR8840397), and SU-1 (Linz et al. 2014) (NCBI SRA accession number SRR8840397), and AS TK-83, TK-84 and TK-85 (Watarai et al. 2019) (NCBI SRA accession numbers SAMD00154502, SAMD00154503, and SAMD00154504, respectively). We first trimmed adapter sequences and reads at low quality positions for all samples using Trim Galore v.0.3.7 using the “stringency 1”, “quality 30” and “length 50” parameters. Quality and adapter trimmed read sets were then mapped against the AP CBS 117618 reference genome (which was downloaded from FungiDB (Stajich et al. 2012)) using the default setting in bwa-mem (Li 2013), converted to sorted bam alignments using samtools v1.14 (Li 2011) and sample names were added with bamaddrg (Watarai et al. 2019).

We performed joint genotyping on the sorted bam alignments using FreeBayes v1.3.5 with the default settings with the exception of setting ploidy to haploid and coverage (-C) to ≥ 20 (Garrison & Marth 2012). We used VcfTools v0.1.14 to filter variants with the following parameters “-remove-indels,” and “-remove-filtered-all,” “-max-missing 1,” “-recode,” and “-recode-INFO-all” (Danecek et al. 2011). This command resulted in a vcf file containing 388,020 sites in which at least one strain had a genotype relative to the AP CBS 117618 reference genome. Variants were annotated using using SNPeff (Cingolani et al. 2012).

### Inferring population structure

We performed several analyses to examine the relationship between AS and AP isolates. Because linked loci can affect population structure inference, we performed further filtering of our SNP VCF file. First, using VCFTools we retained only biallelic sites and sites in which the minor allele frequency was ≥ 0.05. Next, we used PLINK v1.9-beta6.10 with a window size of 50 Kb and a step size of 10 bp to prune SNP pairs where r^2^ ≥ 0.10. This filtering resulted in 598 sites, which were used for population structure analysis. Additionally, we also used a less conservative approach and thinned SNPs such that neighboring SNPs were separated by at least 10 Kb. This filtering resulted in 3,840 SNPs. Both the LD-pruned (P) and non-LD-pruned (NP) datasets were independently used for population structure inference.

First, we used ADMIXTURE to predict the number of ancestral populations (*K*) in our samples (Alexander et al. 2009). We ran 10 replicates for *K* = 1-8 independently on the P and NP datasets. For each analysis we calculated the 10-fold cross-validation error (CV error), in which the lowest value represents the best fit for *K*. For the best values of *K*, we grouped isolates into populations based on their largest membership coefficients (*Q*).

Next, we performed principal components analysis (PCA) on the P and NP datasets in PLINK v1.9-beta6.10 (Bradbury et al. 2007). We plotted eigenvectors for the two eigenvalues that explained the most the highest amount of variance in our dataset.

Finally, we used the alignments of the P and NP datasets to construct maximum likelihood (ML) phylogenetic trees using MEGA-X (Kumar et al. 2018), implementing the generalized time reversible model (GTR) with 100 bootstrap replicates.

### Population Genetics of *A. sojae* and *A. parasiticus*

We used VCFtools to calculate genetic diversity (π) and linkage disequilibrium (*r^2^*) within the AS and AP populations from the full vcf file. π was calculated for each individual site that was polymorphic in at least 1 isolate (N=388,020 sites). *r^2^* was calculated in 5kb windows (65,894 windows) with a 500 bp step size.

### Estimating Copy Number Variation

For each isolate, we estimated integer copy number (CN) for each gene in the reference AP CBS 117618 genome using control-FREEC with the following parameters: window = 1000, step = 200, telocentromeric = 0, minExpectedGC = 0.33, and maxExpectedGC = 0.63 (Steenwyk et al. 2016). Next, we used the BEDtools v2.30.0 (Quinlan & Hall 2010) “intersect” function to identify CNVs (from the control-freec “CNVs” file output) that entirely overlapped gene coordinates gleaned from the gff file. Despite several attempts to run control-freec on AS 3351 and 5597, we opted to remove these samples from the CNV analysis because they contained abnormal copy number profiles and read mapping patterns compared to the other isolates.

Next, we calculated V_ST_, to identify genes with divergent copy number profiles between the AS and AP populations. V_ST_ is analogous to F_ST_, and considers how multiallelic genotype data (such as CNVs) is partitioned between and within population (Rinker et al. 2019). V_ST_ ranges from 0 (no differentiation between groups) to 1 (complete differentiation between groups). V_ST_ was calculated as follows:

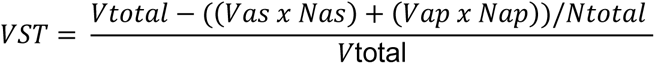

where V = variance, N = population size, “as” = *A. sojae*, and “ap” = *A. parasiticus*. We set a conservative cutoff of V_ST_ ≥ 0.50 to signify genes with differentiated CN patterns between the AS and AP populations.

### Analysis of nonsense mutations fixed in *A. sojae*

We used SnpEff to annotate and predict the function of SNPs that were fixed in AS and had different genotypes in all AP isolates (including AP 1032, 1039, and 2039). We focused on variants that were annotated as nonsense mutations because of their likely detrimental impact on protein function.

### Secondary metabolite gene functional enrichment

To examine the enrichment of genes involved in secondary metabolite encoding gene (SecMet) clusters, we predicted SecMet clusters with antiSMASH v7.0.0. SecMet gene IDs were extracted from the GenBank output file. Enrichment of SecMet genes in different subsets of genes (*e.g.* genes with CNV and genes containing nonsense mutations) were compared against the background using a Fisher’s exact test in JMP Pro 16.

### Sample preparation for phenotypic measurement of *A. sojae* and *A. parasiticus* isolates

Conidia stocks stored in the -80°C freezer were thawed and grown on PDA overnight at 30°C. Plates were then flooded with 50% PDB + 49.9% glycerol + 0.1% tween, and gently scraped with a sterile loop. 10 μL of the conidia suspension was plated onto fresh PDA and this was repeated two additional times. After the third passage, conidia were collected in 1.5 mL 50% PDB + 49.9% glycerol + 0.1% tween and conidia were quantified on a hemocytometer and normalized to 5 × 10^5^ conidia per mL.

### Measuring fungal growth

For each isolate, we measured growth rate on PDA, soy agar (5:1 soy flour to agar ratio), and rice agar (5:1 rice flour to agar ratio). The soy and rice agar mediums represent laboratory “miso” and “sake” fermentation models. For growth rate measurements, 5 × 10^5^ conidia were inoculated onto the center of the plate, incubated at 30°C and automated colony diameter measurements (mm) were collected using the Interscience Scan 1200 at 48 h and 72 h. Three biological replicates were conducted for each isolate and the average colony diameter was used.

### Minimum inhibitory concentration of salt

We measured the minimum inhibitory concentration (MIC) of sodium chloride (NaCl) in miso media for all isolates with NaCl concentrations of 0% to 22% in increments of 2%. MIC values were determined as the first NaCl concentration that visibly lacked growth. 5 x 10^5^ conidia were inoculated into each well of the plate and incubated at 30°C for 48 h and 72 h, at which time each well was individually examined for fungal growth.

### Quantifying protease activity

Protease activity was measured for all isolates in triplicate. 5 x 10^5^ conidia were inoculated into sterile 50ml tubes with 150% hydrated soymeal at 30°C for 48 hours, at which time the Pierce Fluorescent Protease Assay Kit (Thermo Scientific) was used to quantify protease activity following the manufacturer’s instructions. Fluorescence values were measured using a SpectraMax i3 fluorescence microplate reader.

### Aflatoxin Quantification

Isolates were cultured into slant cultures consisting of 2 ml PD broth in 10 ml glass test tubes with ∼10^5^ conidia, inoculated with a loop. The tubes were put in a rack at a 45 ° angle and placed in an incubator at 30°C for 7 days. After 7 days of incubation, aflatoxins were extracted via organic solvents. Chloroform was added and mixed in an amount of 1 mL into the 2 mL of slant culture, respectively. Thereafter, 0.5 mL of the organic solvent portion was extracted after centrifuge at 5,000 g and the organic solvent was evaporated in air. Then, 0.5 mL of HPLC mobile phase (H2O: CH3OH: CH3CN = 50:40:10) was added to the extracted aflatoxin to prepare a sample and filter with 0.45 m filtration unit, which was evaluated by HPLC analysis.

## Results

### *A. sojae* isolates comprise a distinct population

We used two panels of SNPs to infer the population structure of AS and AP. First, we used a subset of SNPs for which SNPs in linkage disequilibrium (*r^2^* > 0.10) were removed, as markers in LD can distort population structure inference. This dataset consisted of 598 SNPs. Second, we used a less conservative approach, and required that SNP sites were separated by a minimum of 10 Kb. This dataset consisted of 3,840 SNPs. These datasets are referred to as “pruned” (P) and “non-pruned” (NP), respectively.

First, we inferred the phylogenetic relationship of the isolates by constructing a maximum likelihood phylogenetic tree (**Figure 1A**, **Figure S1**). This analysis shows that AS samples form a single clade with relatively little genetic variation (exhibited by short branch lengths), while AP samples cluster into several clades. Next, we investigated population structure using the model-based approach implemented in Admixture (Alexander et al. 2009). Cross-validation (CV) error was estimated for each *K* from *K* = 1–8 to estimate the most likely population number, with the lowest CV-error value representing the most likely population number. *K*=2 and *K*=7 displayed the lowest CV-error values, for the P dataset, while *K*=2 and *K*=6 showed the lowest CV-error values for the NP dataset (**Figure 1B**, **Figure S2**). When *K*=2, we observe that all AS isolates are part of the same population, with all samples having membership coefficients (*Q*) of 1 (**Figure 1B**). Interestingly, at *K*=2 AP 1032 and 1039 had minor contributions of the AS population (*Q* = 0.33 and 0.32 for AP 1032 and 1039, respectively) (**Figure 1B**). When *K*=6 in the P or *K*=7 in the NP datasets, AS isolates structured into 2 (NP) or 3 (P) populations, and AP samples structured into 4 populations (AP 1039 and 1032 as a population, AP 2642 as a population, AP 2039, 2040 and CBS 117618 as a population, and AP SU-1, 147, 255 and 81 as a population) (**Figure 1B**, **Figure S2**). Finally, we performed PCA to visualize population structure. In both analyses all AS isolates grouped into a single cluster, AP 1039 and 1032 grouped into a cluster, AP 2642 grouped into its own cluster, and the remaining AP samples grouped into a distinct cluster (**Figure 1C**, **Figure S3**). Collectively, these results provide evidence that AS isolates represent a distinct population when compared with AP isolates.

**Figure 1.**
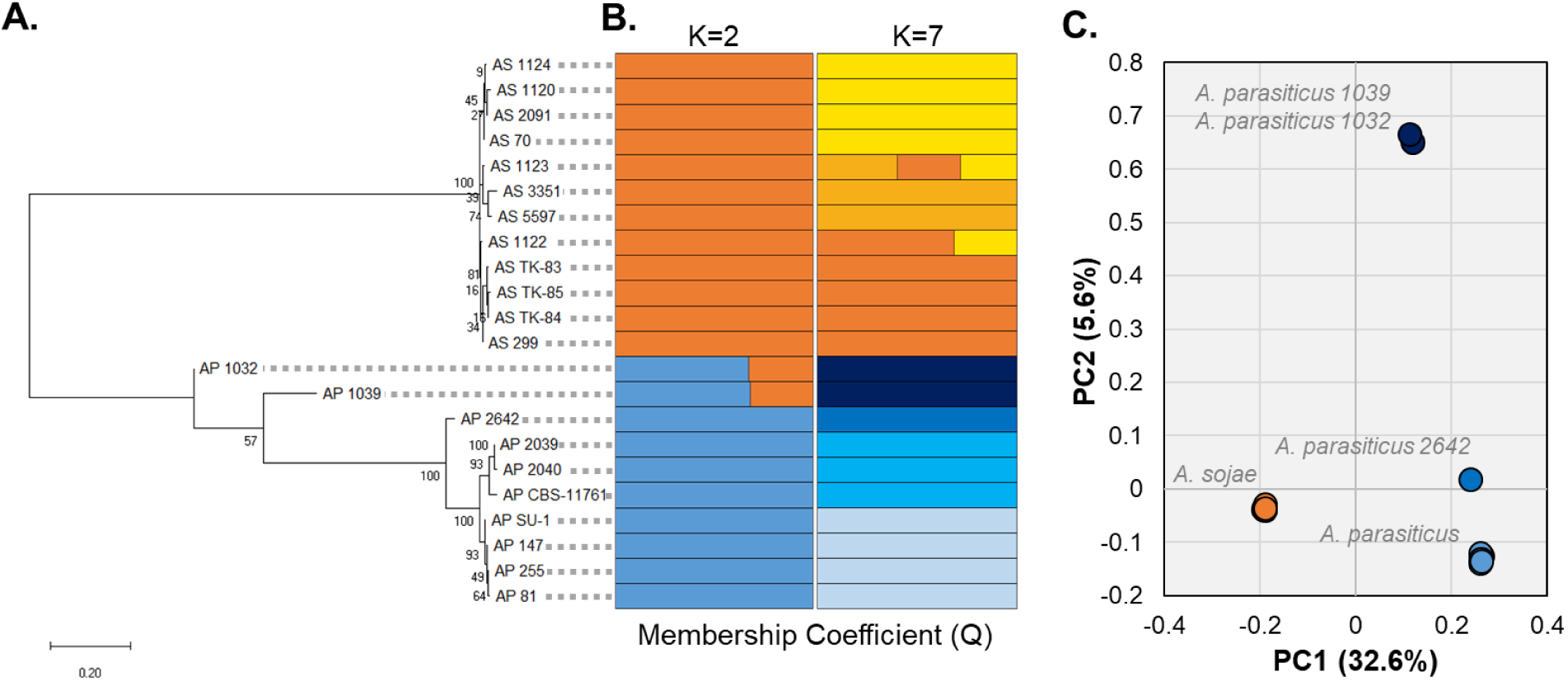
*A. sojae* strains are genetically distinct from *A. parasiticus*. Population structure of *A. sojae* and *A. parasiticus* samples inferred from phylogenetic (A) ADMIXTURE (B) and principal components analysis (C). AS = *A. sojae* (orange) and AP = *A. parasiticus* (blue). For the phylogentic analysis (A) a maximum likelihood tree was constructed and values represent bootstrap support. For ADMIXTURE analysis, membership coefficients are displayed when *K*=2 and *K*=7, as these values had the lowest CV-error scores. PCA (C) shows that all *A. sojae* samples cluster together, while *A. parasiticus* samples cluster into three distinct groups. The percent of variance explained by each principal component is shown in parentheses.

### *A. sojae* displays reduced genetic variation

To better understand the population biology of AS and AP, we calculated genetic diversity and linkage disequilibrium (LD) decay in the AS and AP populations. For these and subsequent analyses, we consider *K*=2 as population number, where all *A. sojae* isolates are grouped into a population (AS) and all *A. parasiticus* isolates are grouped into another population (AP). First, we examined the number of sites that were polymorphic in at least one individual in the *A. sojae* and *A. parasiticus* populations from our filtered vcf file containing 388,020 sites. Strikingly, only 0.61% of sites (2,360 of 388,020) were polymorphic in *A. sojae* compared to 89.74% (348,220 of 388,020) in AP (Fisher’s exact test; *p-value* = 0). We also calculated and compared the average nucleotide diversity (*π*) of the 388,020 sites for *A. sojae* and *A. parasiticus*. Similarly, *π* was very low in AS 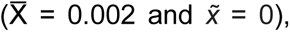 while significantly higher in AP 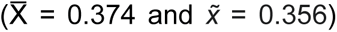 (Wilcoxon signed ranks test; *p-value* = 0) (**Figure 2A**).

**Figure 2.**
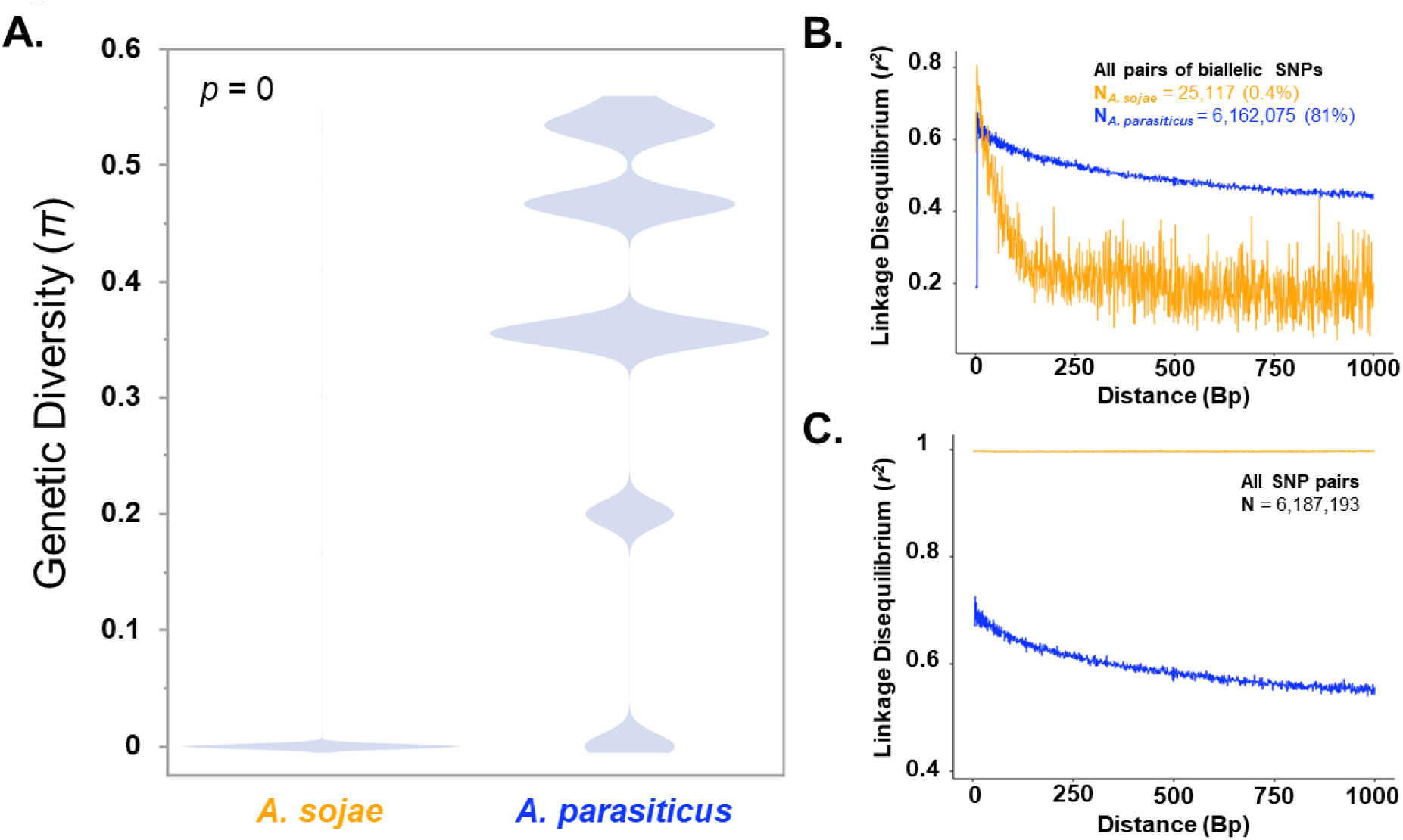
*A. sojae* displays low levels of genetic diversity compared to *A. parasiticus*. (A) Violin plots, displaying the distribution of genetic diversity (π) values across 388,020 sites that were variable in at least one isolate. The p-value is from a Wilcoxon signed ranks test. Linkage disequilibrium decay plots for all pairs of biallelic SNPs (B) and all SNP pairs (C). The X axes represent the physical distance between SNP pairs, and the Y access shows the degree each SNP pair is correlated (*r^2^*). Orange and blue represent *A. sojae* and *A. parasiticus*, respectively.

Next, we calculated LD decay to better understand patterns of recombination in the AS and AP populations. We calculated LD (*r^2^*) within 1 KB bins and analyzed 6,187,193 locus pairs. For AP, 81% of locus pairs were biallelic at both loci, while for AS, only 0.4% of locus pairs were biallelic at both loci (*i.e.* most sites are fixed), owing to the low levels of genetic diversity observed across these isolates (**Figure 2A, B)**. In AP, linkage disequilibrium decayed as a result of physical distance (**Figure 2B, C**). For the small subset of locus pairs that were biallelic in AS, LD decayed with respect to physical distance, however, when all sites were analyzed AS was in almost complete LD (**Figure 2C**). Taken together, these results suggest AS has very low levels of genetic variation (**Figure 2A, C**) but may show patterns of recombination within closely related strains (**Figure 2B**).

### Gene copy number differences between *A. sojae* and *A. parasiticus*

Differences in gene copy number (CN) can be adaptive in closely related populations (Rinker et al. 2019; Steenwyk et al. 2016). Thus, we estimated gene copy number for each of the 13,752 genes with respect to the reference AP CBS 11768 genome and identified genes with different CN profiles across the AS and AP populations. Overall, we found that the number of gene gains (*i.e.* genes with CN ≥ 1) was similar between AS and AP (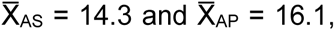 t-test, *p-value* = 0.74), while the number of gene absences (*i.e.* genes with CN = 0) was significantly greater in AS than AP (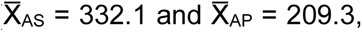 t-test, *p-value* = 0.008) (**Figure S4**). Next, we identified genes for which CN profiles were highly differentiated between AS and AP (*i.e.* genes in which V_ST_ ≥ 0.50). We identified 180 genes displaying differentiated CN profiles between the AS and AP populations (**Table S1**). 162 of these genes had a higher copy number in AP, and represented instances where AP isolates have a CN of 1 and AS isolates have a CN of 0. Nine of the 18 high VST genes with greater CN in AS had an average CN ∼2, including a gene encoding a general substrate transporter, two neighboring genes containing TauD domains, and a predicted allantoinase encoding gene (**Table S1**).

Interestingly, 25.6% of the high V_ST_ genes were annotated as being members of secondary metabolite (SecMet) encoding gene clusters, compared to ∼12% at the whole-genome level (Fisher’s exact text, p-value = 2.12e-5) (**Figure 3**). For example, a gene encoding a general substrate transporter was duplicated in all AS isolates in the aspergillic acid SecMet gene cluster (**Figure 3A**). Additionally, an isoprenoid synthase encoding gene was absent from a predicted terpene encoding gene cluster in all AP isolates with the exception of the AP CBS-117618 reference genome and the two AP isolates (1032 and 1039) most closely related to AS (**Figure 3B)**. Conversely, we also observed large-scale multi-gene deletions in a predicted SecMet gene cluster containing both a nonribosomal peptide synthase and a terpenoid synthase biosynthetic gene and in the predicted Astellolide A gene cluster (**Figures 3C, D**).

**Figure 3.**
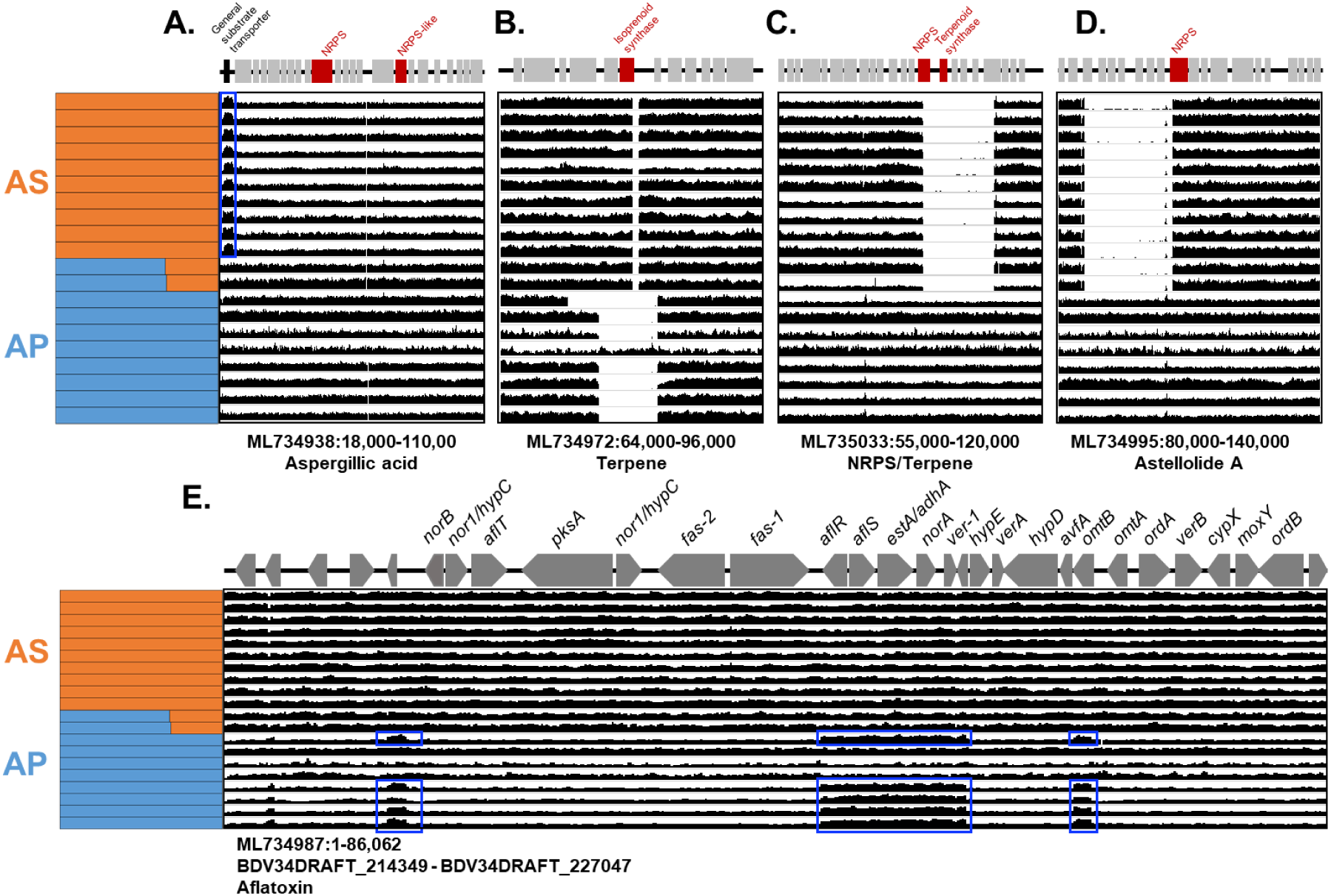
Secondary metabolite encoding gene clusters with regions displaying different copy number profiles between *A. sojae* and *A. parasiticus*. Each row represents an *A. sojae* (orange) or *A. parasiticus* (blue) genome. For each gene cluster (A-E), the scaffold, positions and predicted product are provided. In in A-D, core biosynthetic genes are colored in red, duplicated genes (A) are depicted in black and are also highlighted in blue boxes. For each genome the per site copy number is plotted in black. (E) Gene names are provided for the aflatoxin cluster, with the arrows depicting the direction of transcription. The range of gene identifiers is also listed below the scaffold identifier.

Further, we observed a partial duplication of the aflatoxin gene cluster in four AP isolates (**Figure 3E**). Interestingly, we observed a partial duplication of 1 gene upstream of *norB*, the start of the aflatoxin cluster, and duplications in a six gene region (*aflR, aflS, estA, adhA*, *norA* and ver-1) and *omtB* in AP 2642, SU-1, 147, 255, and 81 (**Figure 3E**). The partial duplication of the aflatoxin cluster in AP has been previously reported (Chang & Yu 2002; Cary et al. 2002).

### Analysis of high impact mutations

Domesticated lineages often accumulate deleterious mutations due to the strength of genetic drift from small population sizes during initial domestication (Mukai et al. 1972). To explore the mutational load in the AS population, we identified mutations that were (i) fixed in AS, (ii) had different genotypes in all AP isolates, and (iii) result in putative nonsense mutations. We focused on nonsense mutations because of their likely impact on protein function. In total, we identified nonsense mutations in 507 protein coding genes. Eighty-one of these genes (∼16%) were annotated by antiSMASH as being members of a SecMet gene cluster, which was a significantly higher proportion than found in the background genome (Fisher’s exact test, p-value = 0.0165). Of note, we identified nonsense mutations in two key genes in the aflatoxin encoding gene cluster, the biosynthetic gene *pksA* (BDV34DRAFT_127106; 5544C>A) and the transcriptional regulator *aflR* (BDV34DRAFT_127154; 1147C>T) (**Figure S5**). This specific *aflR* mutation has been observed previously (Matsushima, Chang, et al. 2001).

### *A. sojae* and *A. parasiticus* grow similarly across substrates and salt concentrations

We hypothesized that AS would grow at a faster rate on food substrates compared to AP because of AS’s usage in, and potentially specialization to, the food environment. We measured growth rate (colony diameter) of all isolates on PDA, rice media and soy media after 48 h and 72 h. Interestingly, growth rate did not significantly differ between AS *and* AP isolates for any of the media types (Student’s t-test; all p-values > 0.05) (**Figure 4**). Next, we hypothesized that variance in growth rate would be reduced in AS to potentially promote uniform fermentation between isolates. We used an F-test to compare variance between the AS and AP populations. We found that variance did not significantly differ in PDA or rice media, but did significantly differ in soy media (48h: F = 0.23, P = 0.037 and 72h: F=0.20, p-value = 0.028) (**Figure 4D**).

**Figure 4.**
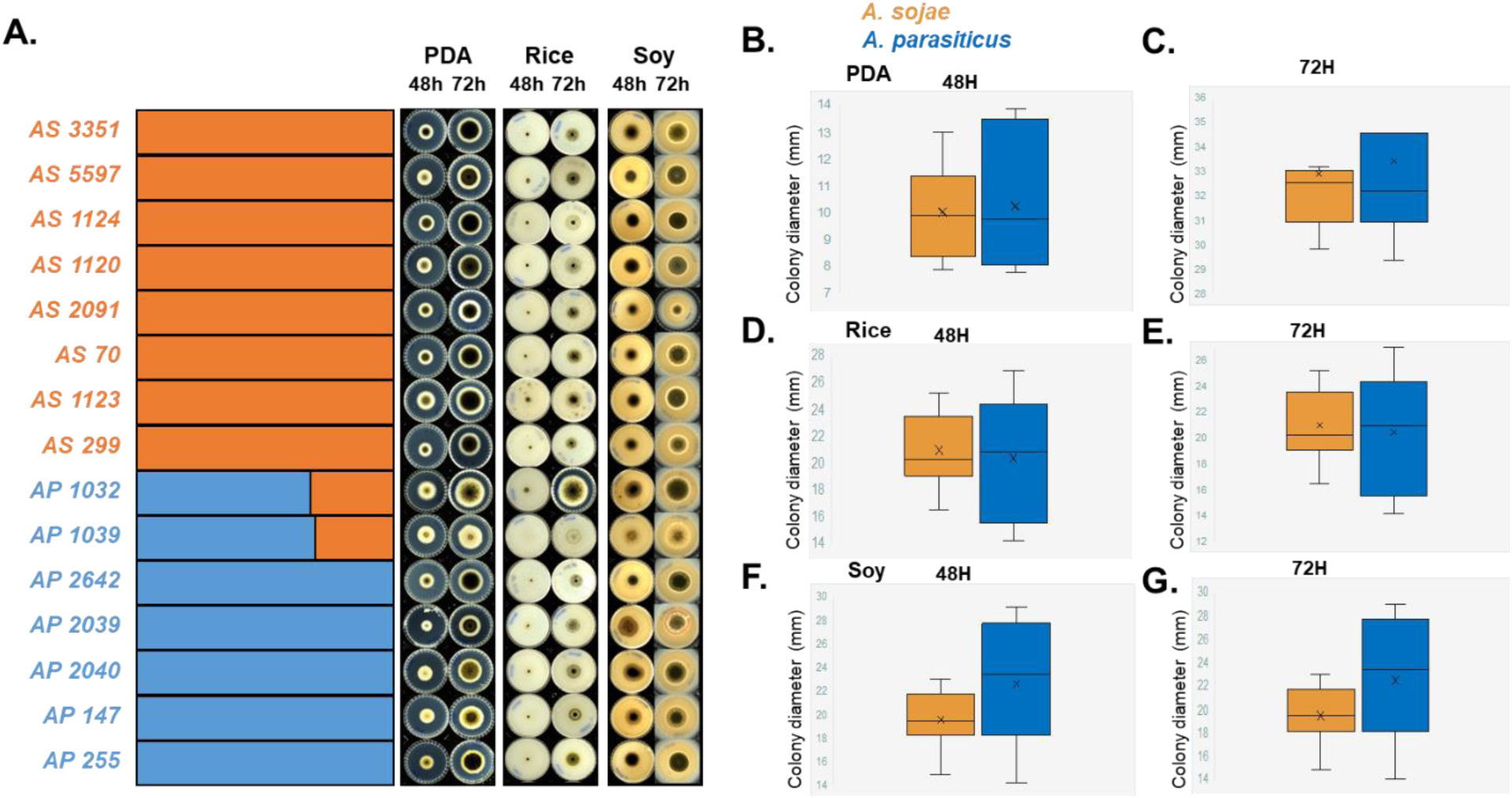
Growth rate on food media does not differ between *A. sojae* and *A. parasiticus*. (A) Strain names are provided with the admixture plot, with *A. sojae* in orange and *A. parasiticus* in blue. Representative images are provided for each strain at 48 h and 72 h during growth on potato dextrose agar, rice media and soy media. Box plots showing colony diameter on potato dextrose agar (B and C), rice (D and E) and soy (F and G) at both time points.

Next, we hypothesized that AS would be more tolerant to high levels of NaCl and would grow at a faster rate in the presence of NaCl because AS is used for miso production (NaCl % in miso is typically 4-6%). Thus, we first measured the MIC of NaCl across all isolates from at 48 h and 72 h. The NaCl MIC did not differ significantly between AS and AP (48h: 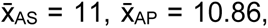 p-value = 0.86 and 72h: 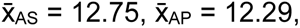 p-value = 0.17) (**Figure S6**). Next, we measured growth rate in soy media enriched with 4% NaCl and 6% NaCl. We did not observe significant differences in growth rate in any comparisons (p-values > 0.05) (**Figure 5**). We again compared variance between AS and AP and found that AS had significantly greater variance in 4% NaCl 48 h (p-value = 0.03), 4% NaCl 72 h (p-value = 4.02e-6), and 6% NaCl 72 h (p-value = 0.006).

**Figure 5.**
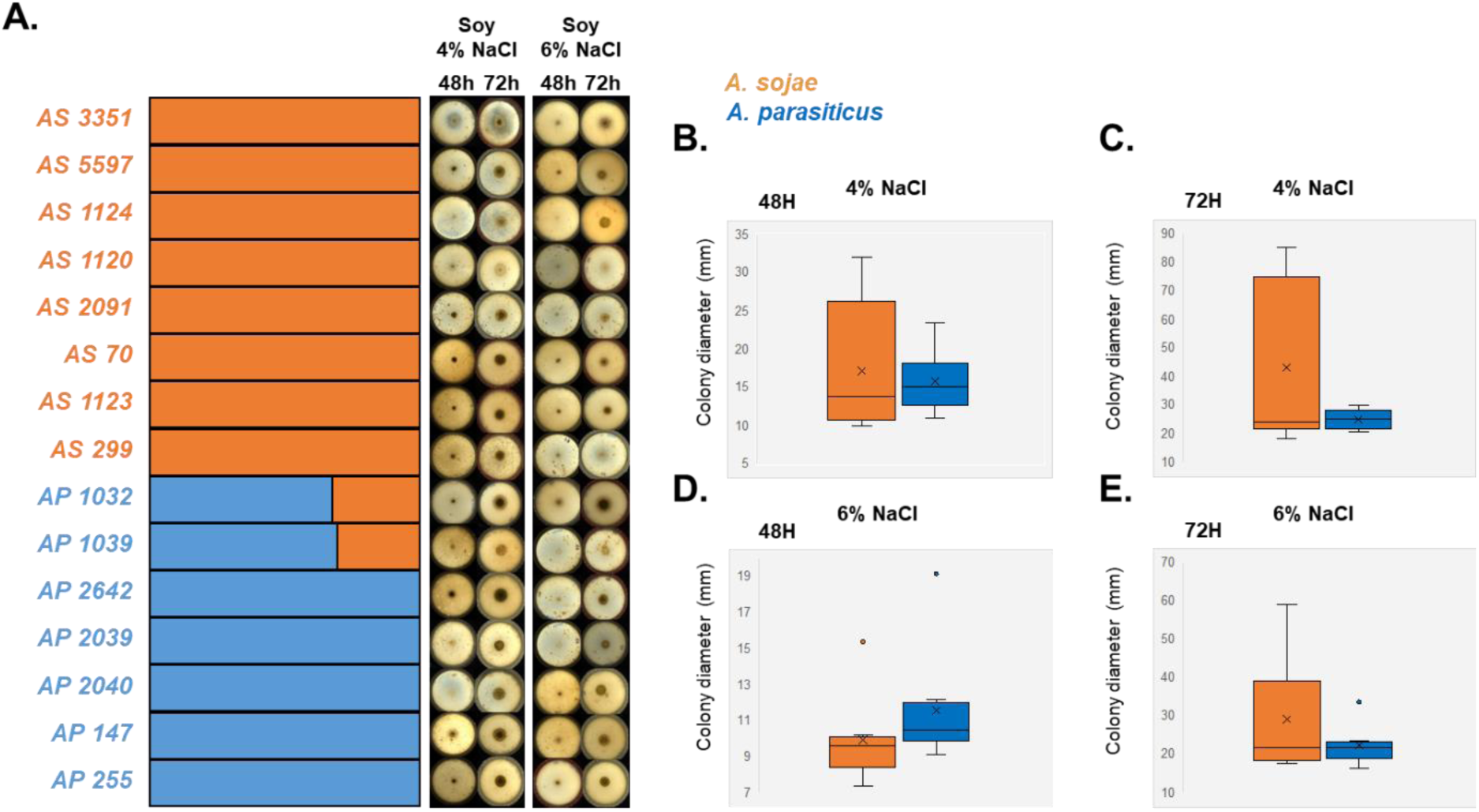
Growth rate with soy media containing salt does not differ between *A. sojae* and *A. parasiticus*. (A) Strain names are provided with the admixture plot, with *A. sojae* in orange and *A. parasiticus* in blue. Representative images are provided for each strain at 48 h and 72 h during growth on soy media with 4% and 6% NaCl. (B-E) Box plots showing colony diameter on soy media for each salt concentration and both time points.

### Protease activity does not differ between *A. sojae* and *A. parasiticus*

AS is used primarily used for soy fermentation (*e.g.* miso and soy sauce) because of AS’s high proteolytic activity (Lim et al. 2019). Thus, we hypothesized that AS would have higher proteolytic activity than AP because of AS’s long-term association with the food environment. Isolates were grown on soymeal for 48 h and protease activity was measured. We did not find a significant difference in protease activity between AS and AP isolates (t-test, p-value = 0.68) (**Figure S7**), or between any isolate pairs (ANOVA, p-value = 0.62).

### *A. sojae* does not produce aflatoxins

Finally, we quantified the production of aflatoxins (AFs) (AFG1, AFG2, AFB1, and AFB2) across 5 AS and 7 AP isolates. No AS isolates produced AF, while the two AP isolates with evidence of ancestry with AS (AP 1032 and 1039) produced AFG2 and AFB1 (**Figure 6**). Additionally, while AP 2039 and 2040 are closely related (Figure 1), AP 2039 did not produce AF while AP 2040 produced AFG1 and AFB1. Lastly, AP 147 produced AFG1 and AFB1 and AP 255 produced AFB1 and AFB2. Aflatoxin production was not correlated with strains possessing the partial duplication of the aflatoxin cluster (**Figure 3E**).

**Figure 6.**
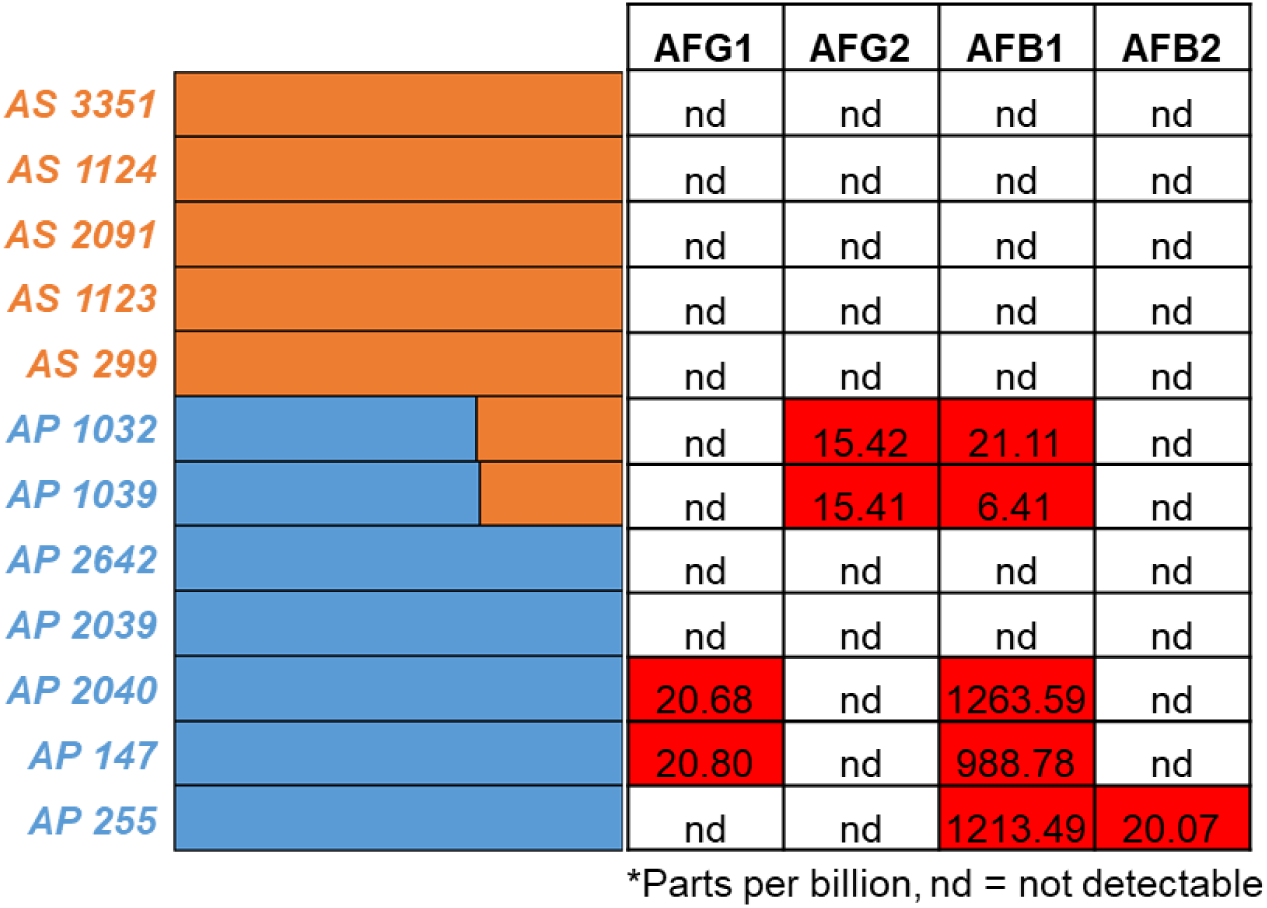
*A. sojae* does not produce aflatoxin. Strain names are provided with the admixture plot, with *A. sojae* in orange and *A. parasiticus* in blue. Aflatoxin G1, G2, B1 and B2 were measured after 7 days of incubation in potato dextrose broth at 30°C. nd = not detectable, while numbered values represent parts per billion. Red boxes indicate aflatoxin production.

## Discussion

We analyzed the genomes and food fermentation characteristics of AS and AP strains to better understand their relationship and their phenotypic differences. Our population genomic analysis revealed that AS is a distinct population from AP (**Figure 1**), and that AS exhibits very low levels of genetic variation (**Figures 1 and 2**). These findings suggest that AS likely evolved from a clonal expansion of a single strain. Interestingly, although polymorphic sites were extremely rare in AS, linkage disequilibrium did decay as a function of physical distance (**Figure 2B**), suggesting that recombination may occur between AS strains. These observations are consistent with other domesticated filamentous molds. For instance, the non-Roquefort blue cheese population of *P. roquefortii* and the soft cheese mold *P. camemberti* display low levels of genetic variation and evidence of recent clonal expansions (Dumas et al. 2020; Ropars et al. 2020). Additionally, *A. oryzae* isolates cluster into at least 8 distinct groups, with each group displaying relatively low levels of genetic variation (Watarai et al. 2019; Chacón-Vargas et al. 2021; Gibbons et al. 2012), again indicating the potential clonal expansion of single strains. In agreement with our results, a recent analysis of seven AS and AP genomes also observed population differentiation between AS and AP and low levels of genetic diversity in AS (Chang & Hua 2023).

Fungal SecMets often function as defense chemicals to fend off competitors, and biosynthesizing these compounds is energetically costly (Keller 2019). We observed widespread restructuring of secondary metabolism in AS compared to AP. For instance, AS strains contained many likely function altering mutations (*i.e.* deletions and nonsense mutations) in several key genes in SecMet encoding clusters. Many of these mutations could conceivably lead to a loss of SecMet production. For instance, AS contains a ∼850 bp deletion overlapping the isoprenoid synthase encoding gene in a terpene encoding gene cluster, deletion of a 7 gene region in the Astellolide A encoding cluster (**Figure 3**), and nonsense mutations resulting in truncated proteins in the aflatoxin encoding cluster genes *pksA* (the core biosynthetic gene) and *aflR* (the master regulatory) (**Figure S5**). The *aflR* premature stop codon mutation in AS strains has been characterized (Chang et al. 2007), and a mutant lacking the AP functional copy, but expressing the AS copy was unable to produce versicolorin A, an aflatoxin intermediate (Takahashi et al. 2002). Other chimeric constructs revealed that, while the AS *aflR* promoter and 5’ region are functional, constructs containing the AS truncated *aflR* were unable to produce aflatoxin (Chang et al. 1999). Additionally, the AS allele of *pksA* contains a premature stop codon resulting in a truncated protein lacking the 3’ thioesterase domain. The PksA protein forms the polyketide backbone from a C6 fatty acid starter unit, and the thioesterase domain is required to hydrolyze the final polyketide chain from the acyl carrier protein domain (Chang et al. 2007). Interestingly, these results indicate that gene clusters encoding SecMets, including aflatoxin, have been accumulating loss-of-function mutations. Unsurprisingly, AS strains did not produce aflatoxin, while 5 of the 7 AP strains, including the closely related AP 1032 and 1039, produced aflatoxin (**Figure 6**).

The loss or reduction of SecMet production is a hallmark of filamentous molds used in fermented food production (Gibbons & Rinker 2015; Steensels et al. 2019). For instance, *A. oryzae* has accumulated a number of mutations in the aflatoxin encoding gene cluster that similarly render *A. oryzae* incapable of producing this toxin (Gibbons et al. 2012; Tominaga et al. 2006; Chang et al. 2005). In *P. camemberti var. “caseifulvum”* strains isolated from non-camemberti white cheeses, a 2 bp deletion is present in the cyclopiazonic acid (CPA) regulatory gene which culminates in a premature stop codon and loss of CPA production (Ropars et al. 2020). Additionally, *P. roqueforti* and *Aspergillus kawachii* (used in the brewing of shochu), have deletions in SecMet clusters that render these species unable to produce mycophenolic acid and ochratoxin A, respectively (Gillot et al. 2017; Futagami et al. 2011). The restructuring of secondary metabolism could be the result of artificial selection acting to maintain the microbial community in the fermented foods, as SecMets are often defense compounds with antimicrobial properties, or for the reallocation of energy from secondary metabolism to primary metabolism (Gibbons 2019).

To produce miso, AS spores are inoculated into cooled steamed rice and fermented for 2 days at 30°C. The fermented rice is then mixed with cooked and mashed soybeans and salt, and fermented in containers for up to two years (Allwood et al. 2021). Thus, we hypothesized that AS would outperform AP in terms of growth rate on food substrates and tolerance to NaCl because of AS’s usage in rice and soy fermentation. However, other than the absence of aflatoxin production in AS (**Figure 6**), we observed no major differences in growth rate on rice or soy in the absence or presence of salt (**Figures 4, 5**), tolerance to salt (**Figure S6**), or protease activity (**Figure S7**). While these results did not support our hypothesis, our strains were grown at a much shorter duration than in traditional miso fermentation (*i.e.* 2 or 3 days vs. months), and in the absence of microbial community members found in miso. For instance, amplicon sequencing of miso samples matured for less than six months and greater than six months all showed the presence of *Aspergillus* sp. (Allwood et al. 2023). Thus, phenotypic differences between AS and AP may not be apparent until later stages of fermentation, where AS may still be metabolically active. Additionally, faster growth rate might not be a target of selection, as is the case with the Roquefort population of *P. roqueforti*, in which slow maturation of cheese is preferred to enhance preservation and limit degradation (Dumas et al. 2020). Lastly, the influence of AS and AP on the microbes in the fermentation community is unknown. The production of SecMets by AP could inhibit the growth of the bacteria and yeasts found in the food microbial community. For instance, aflatoxin shows some antibacterial and antiyeast effects (Ali-Vehmas et al. 1998; Keller-Seitz et al. 2004), which could alter the miso microbial community and negatively impact the sensory attributes of miso.

Collectively, we analyzed the genomes and phenotypes of various AS and AP strains. We found distinct genetic differentiation between AS from AP, evidence of a recent clonal expansion in AS, and the loss of toxin production in AS, all of which resemble patterns in other domesticated filamentous fungi. Thus, we hypothesize that AS was domesticated relatively recently from a single strain. Deeper sampling and further phenotypic characterization of AS and AP strains will be required to further shed light on the impacts of domestication on AS.

## Acknowledgments

This work was funded through the National Science Foundation Grant 1942681 to JGG, which also supports KLA. KLA is also supported by a Spaulding Smith Fellowship from the University of Massachusetts Amherst. The work at UW-Madison was support by the National Institute of Food and Agriculture, United States Department of Agriculture, Hatch project 7000326 to JHY.

## Data Availability Statement

Raw whole-genome Illumina data for all isolates is available through the NCBI Sequence Read Archive through BioProject accession number PRJNA911610.

