## Supplementary material for "Population genomics of *Aspergillus sojae* is shaped by the food environment": Figure S1

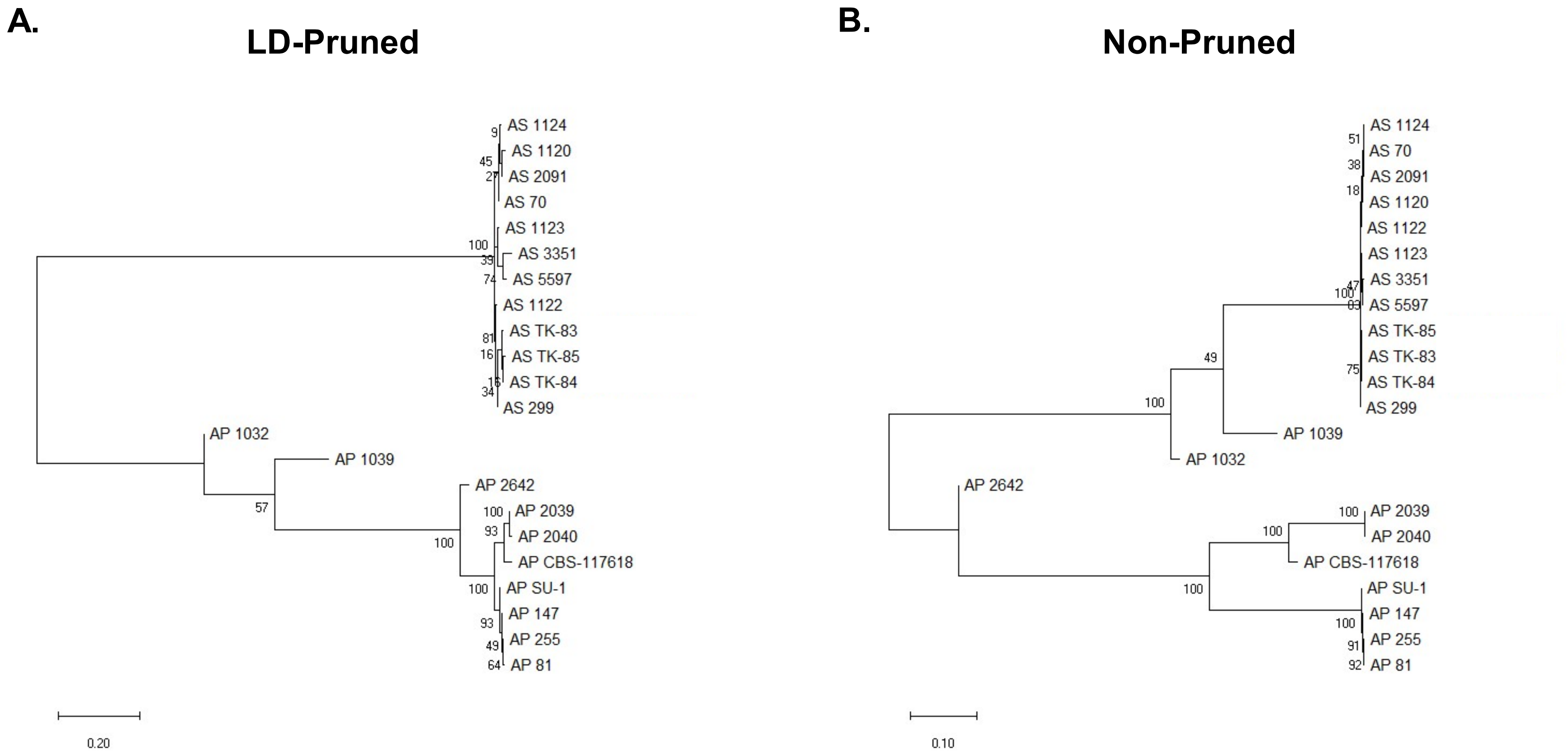


**Figure S1. Phylogenetic relationship of *A. sojae* and *A. parasiticus* isolates.** Maximum likelihood tree were constructed using the LD-pruned (A) and non-pruned (B) SNP datasets using the GTR model. Trees are rooted by the midpoint. Values represent bootstrap support for 100 bootstrap replicates.
