## Supplementary material for "Population genomics of *Aspergillus sojae* is shaped by the food environment": Figure S2

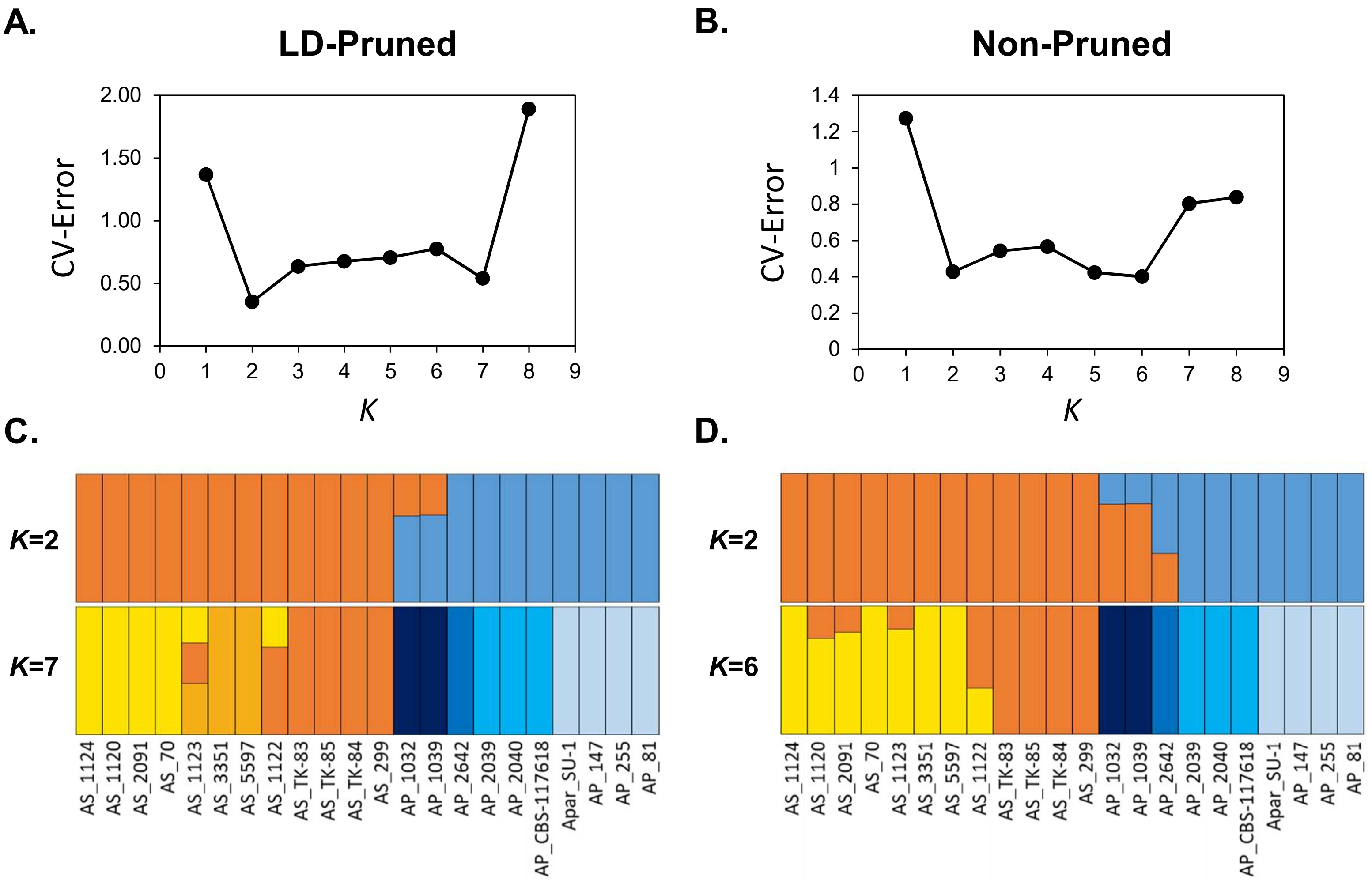


**Figure S2. Population structure of *A. sojae* and *A. parasiticus* isolates.** CV-error plots for the LD-pruned (A) and non-pruned (B) SNP datasets. Admixture plots displaying membership coefficients (*Q*) for *K* values with the lowest CV-error values in the LD-pruned (A) and non-pruned (B) datasets.
