## Supplementary material for "Population genomics of *Aspergillus sojae* is shaped by the food environment": Figure S3

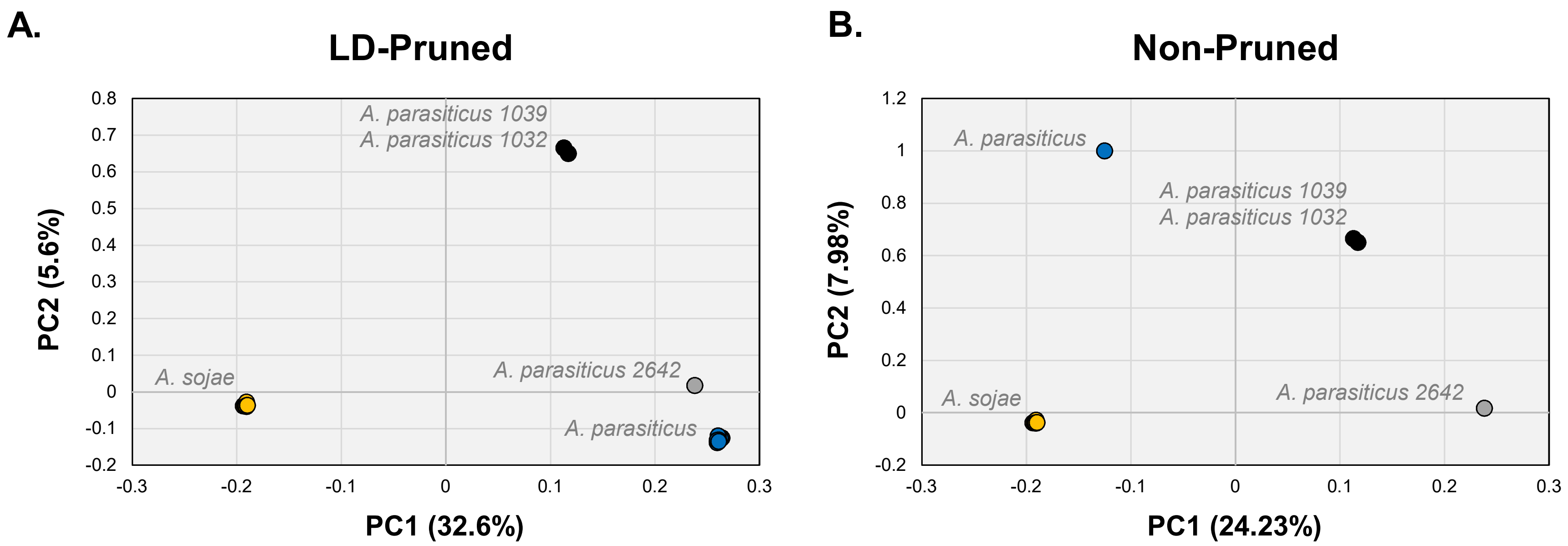


**Figure S3. Principal components analysis (PCA) of *A. sojae* and *A. parasiticus* isolates.** PCA analysis was conducted with SNP data for the LD-pruned (A) and non-pruned (B) datasets. The two PCs explaining the most variance are plotted as PC1 and PC2 and the percentage of variance explained is reported. In both cases, AS samples cluster into one group, while AP samples cluster into three distinct groups.
