## Supplementary material for "Population genomics of *Aspergillus sojae* is shaped by the food environment": Figure S4

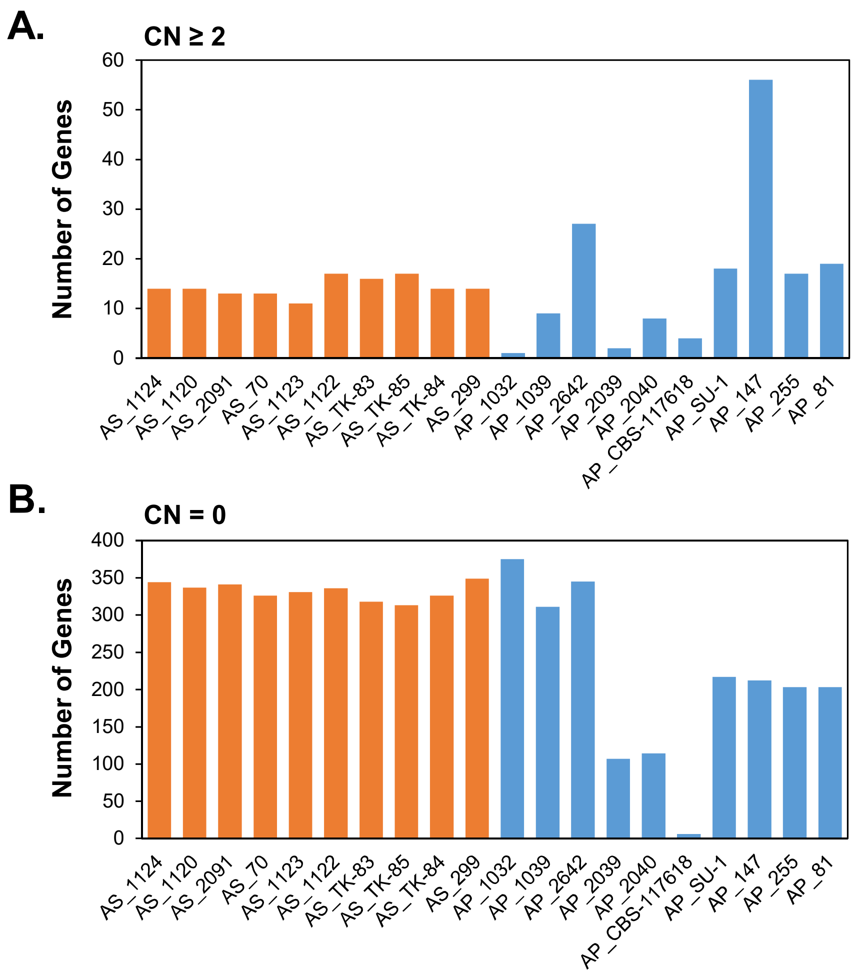


**Figure S4. Genes displaying copy number variation across *A. sojae* and *A. parasiticus* isolates.** Gene gains (A) and gene absences (B) are plotted for each isolate. *A. sojae* samples are in orange and *A. parasiticus* samples are in blue.
