## Supplementary material for "Population genomics of *Aspergillus sojae* is shaped by the food environment": Figure S5

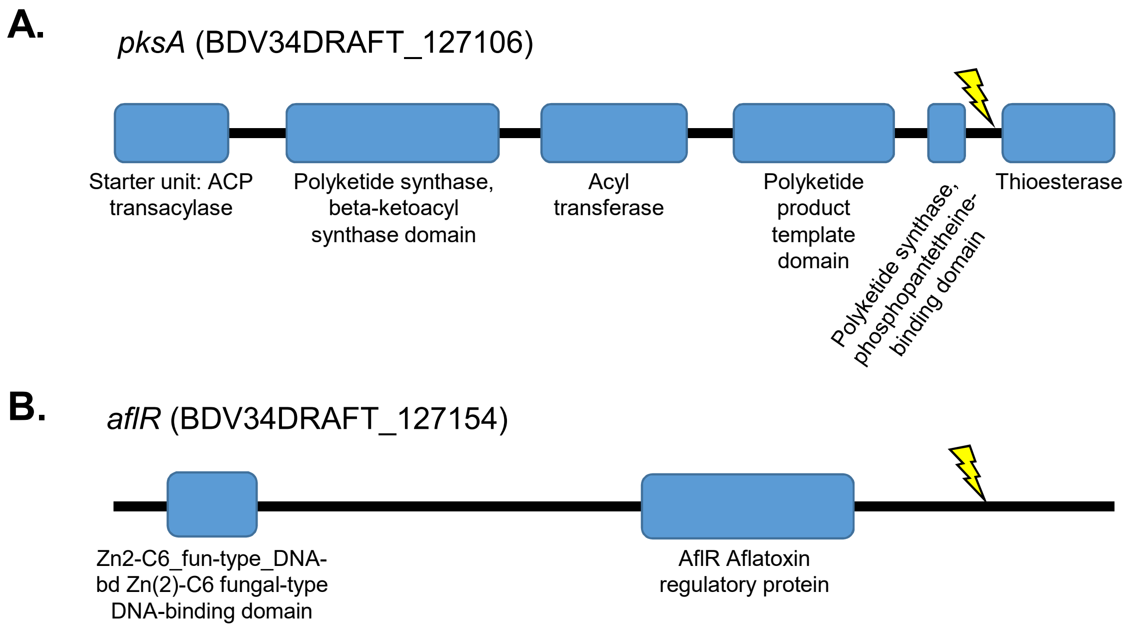


**Figure S5. Schematic of *A. sojae* mutations in aflatoxin encoding cluster genes.** Schematic of *pskA* (A) and *aflR* (B) proteins with ellipses representing InterPro domains. Gene IDs relative to the A. parasiticus CBS-117618 reference genome are in parentheses. The yellow symbol shows the location of nonsense mutations.
