## Supplementary material for "Population genomics of *Aspergillus sojae* is shaped by the food environment": Figure S6

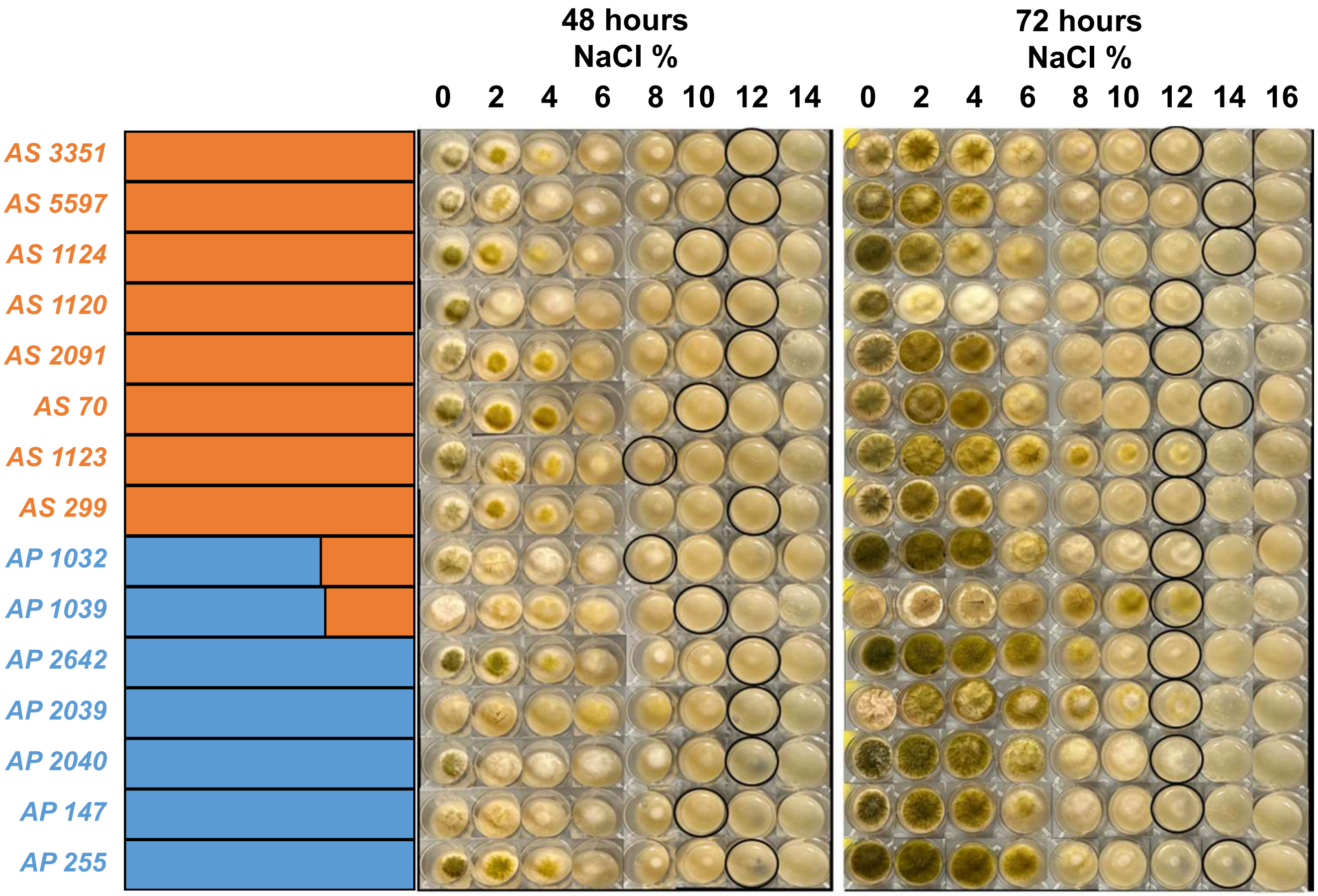


**Figure S6. Minimum inhibitory concentration of NaCl on *A. sojae* and *A. parasiticus*.** Strain names are provided with the admixture plot, with *A. sojae* in orange and *A. parasiticus* in blue. Representative images are provided for each strain at 48 h and 72 h during growth on soy media supplemented with 0, 2, 4, 6, 8, 10, 12, 14 and 16% NaCl. Black circles indicate the highest concentration with observable growth.
