## Supplementary material for "Population genomics of *Aspergillus sojae* is shaped by the food environment": Figure S7

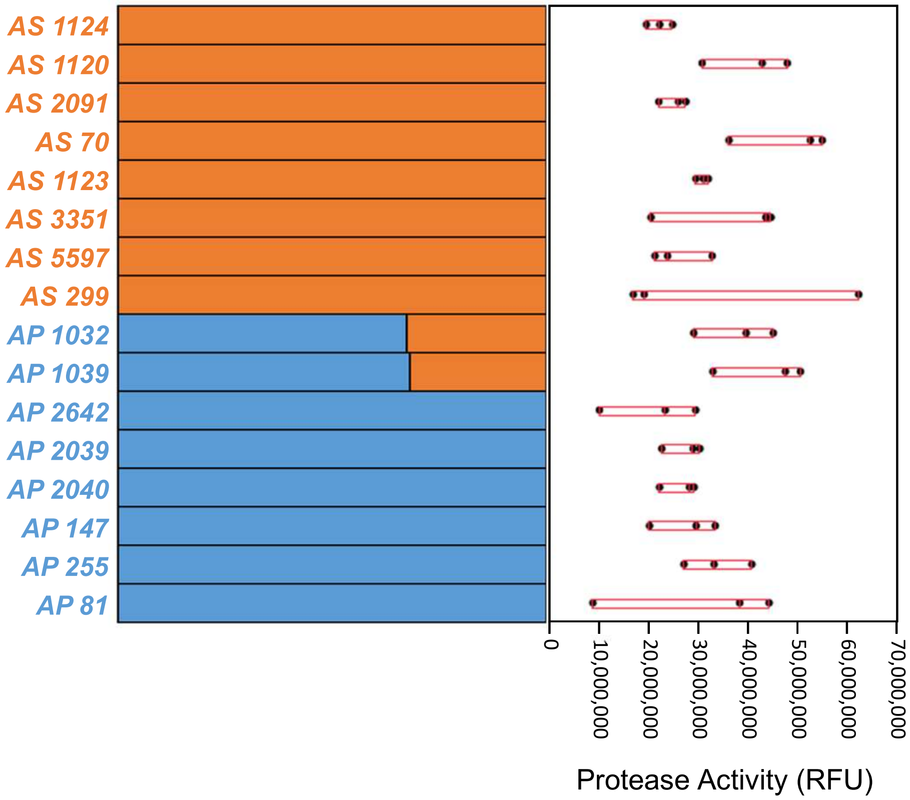


**Figure S7. Protease activity for *A. sojae* and *A. parasiticus* isolates.** Strain names are provided with the admixture plot, with *A. sojae* in orange and *A. parasiticus* in blue. Protease activity was measured for all isolates in triplicate after 48 h of growth at 30°C on 150% hydrated soymeal. Protease activity was measured in terms of relative fluorescence units (RFUs) (X axis).
