## Supplementary material for "Population genomics of *Aspergillus sojae* is shaped by the food environment": Table S1

**Table S1. Genes with highly differentiated copy number between AS and AP isolates.**

| **Scaff** | **gene_start** | **gene_stop** | **gene_ID** | **gene anno** | **Av AS** | **Av AP** | **VST** |
| --- | --- | --- | --- | --- | --- | --- | --- |
| ML734936 | 437368 | 437858 | BDV34DRAFT_218708 | Glyco_hydro_38C domain-containing protein | 0 | 0.8 | 0.65 |
| ML734936 | 726193 | 726967 | BDV34DRAFT_231871 | N-acetyltransferase domain-containing protein | 1 | 0.3 | 0.51 |
| ML734937 | 156349 | 157513 | BDV34DRAFT_61776 | BTB domain-containing protein | 0 | 0.9 | 0.81 |
| ML734937 | 264882 | 268358 | BDV34DRAFT_219087 | CHAT domain-containing protein | 0 | 0.7 | 0.51 |
| ML734937 | 268781 | 269773 | BDV34DRAFT_219088 | Fucose-specific lectin | 0 | 0.7 | 0.51 |
| ML734937 | 269884 | 273909 | BDV34DRAFT_233882 | AAA domain-containing protein | 0 | 0.7 | 0.51 |
| ML734937 | 463614 | 464783 | BDV34DRAFT_219161 | nucleoside phosphorylase domain-containing protein | 0 | 0.9 | 0.81 |
| ML734938 | 19159 | 21033 | BDV34DRAFT_234721 | general substrate transporter | 2 | 1 | 1.00 |
| ML734938 | 247433 | 249471 | BDV34DRAFT_86146 | ANK_REP_REGION domain-containing protein | 0 | 0.8 | 0.65 |
| ML734938 | 249489 | 252728 | BDV34DRAFT_184831 | C2H2-type domain-containing protein | 0 | 0.8 | 0.65 |
| ML734938 | 336721 | 337257 | BDV34DRAFT_219493 | Protein kinase domain-containing protein | 0 | 0.9 | 0.81 |
| ML734938 | 471366 | 474711 | BDV34DRAFT_234813 | Carrier domain-containing protein | 0 | 0.9 | 0.81 |
| ML734938 | 921287 | 921997 | BDV34DRAFT_219707 | Retrotransposon protein | 1.9 | 1 | 0.81 |
| ML734939 | 152488 | 152799 | BDV34DRAFT_219765 | Transposase | 0 | 0.9 | 0.81 |
| ML734939 | 153411 | 154785 | BDV34DRAFT_185265 | FAD-binding PCMH-type domain-containing protein | 0 | 0.9 | 0.81 |
| ML734939 | 205562 | 206628 | BDV34DRAFT_185296 | Translationally-controlled tumor protein | 0 | 0.8 | 0.65 |
| ML734939 | 336885 | 339953 | BDV34DRAFT_235452 | KR domain-containing protein | 0 | 0.8 | 0.65 |
| ML734940 | 287915 | 289804 | BDV34DRAFT_124419 | Integral membrane protein | 0 | 0.9 | 0.81 |
| ML734940 | 289967 | 290293 | BDV34DRAFT_220122 | Beta-lactamase domain-containing protein | 0 | 0.9 | 0.81 |
| ML734940 | 424344 | 425310 | BDV34DRAFT_220172 | DUF2326 domain-containing protein | 1 | 0.3 | 0.51 |
| ML734940 | 425889 | 427534 | BDV34DRAFT_220173 | P-loop containing nucleoside triphosphate hydrolase protein | 1 | 0.3 | 0.51 |
| ML734940 | 524984 | 526216 | BDV34DRAFT_220211 | MBOAT_2 domain-containing protein | 0 | 0.9 | 0.81 |
| ML734941 | 357372 | 357860 | BDV34DRAFT_220390 | Endoribonuclease L-PSP/chorismate mutase-like protein | 0 | 0.8 | 0.65 |
| ML734941 | 358132 | 359307 | BDV34DRAFT_220391 | PARP catalytic domain-containing protein | 0 | 0.8 | 0.65 |
| ML734943 | 614709 | 616847 | BDV34DRAFT_209733 | monooxygenase | 0 | 0.8 | 0.65 |
| ML734946 | 545813 | 547568 | BDV34DRAFT_232308 | ankyrin repeat-containing domain protein | 0 | 1 | 1.00 |
| ML734946 | 572714 | 574648 | BDV34DRAFT_221597 | ANK_REP_REGION domain-containing protein | 0 | 0.9 | 0.81 |
| ML734947 | 296699 | 298046 | BDV34DRAFT_232556 | fido domain-containing protein | 0 | 0.7 | 0.51 |
| ML734949 | 318212 | 318795 | BDV34DRAFT_222146 | Protein kinase domain-containing protein | 1.7 | 1 | 0.51 |
| ML734949 | 377201 | 378307 | BDV34DRAFT_222168 | FAD/NAD(P)-binding domain-containing protein | 0 | 0.9 | 0.81 |
| ML734949 | 379415 | 379843 | BDV34DRAFT_222169 | ankyrin repeat-containing domain protein | 0 | 0.9 | 0.81 |
| ML734949 | 380294 | 381722 | BDV34DRAFT_189736 | Zn(2)-C6 fungal-type domain-containing protein | 0 | 0.9 | 0.81 |
| ML734949 | 382087 | 383738 | BDV34DRAFT_233003 | amidase | 0 | 0.9 | 0.81 |
| ML734949 | 384137 | 385405 | BDV34DRAFT_222174 | MFS general substrate transporter | 0 | 0.9 | 0.81 |
| ML734949 | 544325 | 546246 | BDV34DRAFT_189795 | GSP_synth domain-containing protein | 1.8 | 1 | 0.65 |
| ML734950 | 2567 | 3084 | BDV34DRAFT_222251 | Phosphatidylglycerol/phosphatidylinositol transfer protein | 0 | 0.8 | 0.65 |
| ML734950 | 108943 | 109642 | BDV34DRAFT_222293 | C2H2-type domain-containing protein | 1 | 0.3 | 0.51 |
| ML734951 | 2982 | 4345 | BDV34DRAFT_233279 | TauD domain-containing protein | 2 | 1 | 1.00 |
| ML734951 | 4596 | 6237 | BDV34DRAFT_210805 | TauD domain-containing protein | 2 | 1 | 1.00 |
| ML734951 | 423938 | 424606 | BDV34DRAFT_210920 | O-methyltransferase | 0 | 1 | 1.00 |
| ML734952 | 155043 | 156770 | BDV34DRAFT_222698 | PHD-type domain-containing protein | 0 | 0.7 | 0.51 |
| ML734952 | 160391 | 161380 | BDV34DRAFT_222700 | Uma2 domain-containing protein | 0 | 0.7 | 0.51 |
| ML734952 | 171559 | 172855 | BDV34DRAFT_190636 | Cadherin domain-containing protein | 0 | 0.7 | 0.51 |
| ML734953 | 488522 | 488863 | BDV34DRAFT_222991 | Response regulatory domain-containing protein | 0.1 | 0.9 | 0.62 |
| ML734954 | 331856 | 334074 | BDV34DRAFT_191377 | HSF_DOMAIN domain-containing protein | 0 | 0.7 | 0.51 |
| ML734955 | 36481 | 36864 | BDV34DRAFT_223189 | Secreted protein | 0 | 0.8 | 0.65 |
| ML734955 | 70431 | 71263 | BDV34DRAFT_223201 | Transmembrane protein | 0 | 0.8 | 0.65 |
| ML734955 | 378839 | 379578 | BDV34DRAFT_223316 | Secreted protein | 0 | 0.8 | 0.65 |
| ML734956 | 251228 | 252822 | BDV34DRAFT_211566 | CHAT domain-containing protein | 0 | 0.7 | 0.51 |
| ML734956 | 423885 | 424498 | BDV34DRAFT_223508 | XK-related protein | 0.1 | 0.9 | 0.62 |
| ML734956 | 430767 | 434985 | BDV34DRAFT_192001 | Chitinase | 0 | 0.7 | 0.51 |
| ML734957 | 24880 | 25640 | BDV34DRAFT_192021 | bulb-type lectin domain-containing protein | 0.3 | 1 | 0.51 |
| ML734958 | 102524 | 103427 | BDV34DRAFT_223722 | NmrA domain-containing protein | 0 | 0.9 | 0.81 |
| ML734958 | 285816 | 286423 | BDV34DRAFT_223783 | CVNH domain-containing protein | 0 | 0.7 | 0.51 |
| ML734958 | 329626 | 330991 | BDV34DRAFT_211818 | Integral membrane protein | 0 | 0.9 | 0.81 |
| ML734959 | 243495 | 244826 | BDV34DRAFT_192709 | Ricin B-type lectin domain-containing protein | 0 | 0.7 | 0.51 |
| ML734962 | 142878 | 143536 | BDV34DRAFT_79962 | Tail fiber assembly protein | 0 | 0.8 | 0.65 |
| ML734962 | 273731 | 274420 | BDV34DRAFT_193607 | SDR family NAD(P)-dependent oxidoreductase | 0 | 0.8 | 0.65 |
| ML734962 | 275850 | 276611 | BDV34DRAFT_193608 | unspecified product | 0 | 0.9 | 0.81 |
| ML734962 | 342237 | 343926 | BDV34DRAFT_212276 | APH domain-containing protein | 0 | 1 | 1.00 |
| ML734962 | 357047 | 357710 | BDV34DRAFT_224399 | Secreted protein | 0 | 0.9 | 0.81 |
| ML734962 | 397806 | 400070 | BDV34DRAFT_234589 | actin-like ATPase domain-containing protein | 0 | 0.9 | 0.81 |
| ML734962 | 400325 | 401482 | BDV34DRAFT_224415 | unspecified product | 0 | 0.9 | 0.81 |
| ML734962 | 401942 | 402288 | BDV34DRAFT_212293 | Ubiquitin-like protein 5 | 0 | 0.9 | 0.81 |
| ML734962 | 403445 | 403902 | BDV34DRAFT_224417 | DUF418 domain-containing protein | 0 | 0.9 | 0.81 |
| ML734965 | 107329 | 108616 | BDV34DRAFT_90196 | Hemerythrin domain-containing protein | 0 | 0.8 | 0.65 |
| ML734966 | 132743 | 136392 | BDV34DRAFT_212667 | major facilitator superfamily domain-containing protein | 0 | 1 | 1.00 |
| ML734968 | 42443 | 43862 | BDV34DRAFT_225098 | peptidase S8/S53 domain-containing protein | 0 | 0.8 | 0.65 |
| ML734968 | 45315 | 46987 | BDV34DRAFT_225099 | AA_permease domain-containing protein | 0 | 0.8 | 0.65 |
| ML734968 | 48504 | 49004 | BDV34DRAFT_225100 | FAD_binding_3 domain-containing protein | 0 | 0.8 | 0.65 |
| ML734968 | 50329 | 51589 | BDV34DRAFT_235067 | Similar to | 0 | 0.7 | 0.51 |
| ML734970 | 301452 | 302567 | BDV34DRAFT_225452 | Deacetylase sirtuin-type domain-containing protein | 0 | 1 | 1.00 |
| ML734971 | 173490 | 174574 | BDV34DRAFT_101879 | S-adenosyl-L-methionine-dependent methyltransferase | 0 | 0.7 | 0.51 |
| ML734971 | 174574 | 175307 | BDV34DRAFT_101876 | Secreted protein | 0 | 0.7 | 0.51 |
| ML734971 | 175445 | 176387 | BDV34DRAFT_195825 | DUF362 domain-containing protein | 0 | 0.7 | 0.51 |
| ML734971 | 263948 | 265807 | BDV34DRAFT_195853 | unspecified product | 0 | 0.8 | 0.65 |
| ML734972 | 76746 | 78363 | BDV34DRAFT_213222 | cytochrome P450 | 1 | 0.3 | 0.51 |
| ML734972 | 78661 | 80280 | BDV34DRAFT_195961 | isoprenoid synthase domain-containing protein | 1 | 0.3 | 0.51 |
| ML734974 | 170758 | 172033 | BDV34DRAFT_235379 | C2 domain-containing protein | 0.1 | 0.9 | 0.62 |
| ML734974 | 219545 | 220470 | BDV34DRAFT_196430 | unspecified product | 0 | 0.9 | 0.81 |
| ML734974 | 237055 | 238118 | BDV34DRAFT_225865 | DUF2236 domain-containing protein | 0 | 0.8 | 0.65 |
| ML734975 | 27352 | 29209 | BDV34DRAFT_225899 | MYND-type domain-containing protein | 0 | 0.8 | 0.65 |
| ML734975 | 191528 | 193245 | BDV34DRAFT_196586 | kinase-like protein | 0 | 0.7 | 0.51 |
| ML734976 | 68679 | 72035 | BDV34DRAFT_226022 | Non-specific serine/threonine protein kinase | 0 | 0.7 | 0.51 |
| ML734976 | 89819 | 90880 | BDV34DRAFT_226030 | Neprosin domain-containing protein | 0 | 0.9 | 0.81 |
| ML734976 | 170368 | 172031 | BDV34DRAFT_213580 | transmembrane amino acid transporter protein-domain-containing protein | 0 | 0.8 | 0.65 |
| ML734976 | 173325 | 175824 | BDV34DRAFT_226062 | But2 domain-containing protein | 0 | 0.8 | 0.65 |
| ML734980 | 124450 | 126237 | BDV34DRAFT_197444 | tryptophan RNA-binding attenuator protein-like domain-containing protein | 1 | 0.3 | 0.51 |
| ML734982 | 207562 | 208969 | BDV34DRAFT_197850 | unspecified product | 0 | 0.7 | 0.51 |
| ML734983 | 127736 | 130040 | BDV34DRAFT_235854 | kinase-like domain-containing protein | 0 | 0.9 | 0.81 |
| ML734984 | 40661 | 48585 | BDV34DRAFT_226764 | ketoacyl-synt-domain-containing protein | 0 | 0.7 | 0.51 |
| ML734987 | 110875 | 111312 | BDV34DRAFT_227060 | Secreted protein | 0 | 1 | 1.00 |
| ML734987 | 166610 | 169102 | BDV34DRAFT_127572 | fungal-specific transcription factor domain-containing protein | 0 | 0.7 | 0.51 |
| ML734991 | 131523 | 133544 | BDV34DRAFT_132695 | Mg2+ transporter protein%2C CorA-like/Zinc transport protein ZntB | 0 | 0.8 | 0.65 |
| ML734992 | 39977 | 42129 | BDV34DRAFT_214701 | PcfJ domain-containing protein | 0 | 0.7 | 0.51 |
| ML734995 | 85714 | 88006 | BDV34DRAFT_236594 | cytochrome P450 | 0 | 0.8 | 0.65 |
| ML734995 | 88071 | 88676 | BDV34DRAFT_236596 | HAD-like domain-containing protein | 0 | 0.8 | 0.65 |
| ML734995 | 89154 | 90980 | BDV34DRAFT_236597 | major facilitator superfamily domain-containing protein | 0 | 0.8 | 0.65 |
| ML734995 | 91539 | 93117 | BDV34DRAFT_214918 | transferase | 0 | 0.8 | 0.65 |
| ML734995 | 93711 | 95276 | BDV34DRAFT_236598 | cytochrome P450 | 0 | 0.8 | 0.65 |
| ML734995 | 95673 | 96458 | BDV34DRAFT_236600 | Glucose 1-dehydrogenase | 0 | 0.8 | 0.65 |
| ML734995 | 97615 | 99363 | BDV34DRAFT_236602 | cytochrome P450 | 0 | 0.8 | 0.65 |
| ML734995 | 100224 | 101949 | BDV34DRAFT_199839 | HAD-like domain-containing protein | 0 | 0.8 | 0.65 |
| ML734995 | 102591 | 104339 | BDV34DRAFT_199841 | cytochrome P450 | 0 | 0.8 | 0.65 |
| ML735001 | 51459 | 53099 | BDV34DRAFT_200561 | NADP-dependent oxidoreductase domain-containing protein | 0 | 0.8 | 0.65 |
| ML735001 | 53651 | 54902 | BDV34DRAFT_236811 | BZIP domain-containing protein | 0 | 0.8 | 0.65 |
| ML735001 | 55555 | 57306 | BDV34DRAFT_236813 | FAD-binding PCMH-type domain-containing protein | 0 | 0.8 | 0.65 |
| ML735002 | 156355 | 160038 | BDV34DRAFT_148643 | ankyrin repeat-containing domain protein | 0 | 1 | 1.00 |
| ML735002 | 160596 | 161919 | BDV34DRAFT_148677 | nucleoside phosphorylase domain-containing protein | 0 | 1 | 1.00 |
| ML735002 | 162604 | 163748 | BDV34DRAFT_200742 | DUF1275 domain protein | 0 | 1 | 1.00 |
| ML735002 | 163809 | 165501 | BDV34DRAFT_236881 | general substrate transporter | 0 | 1 | 1.00 |
| ML735004 | 151796 | 153346 | BDV34DRAFT_215502 | cytochrome protein | 0 | 0.8 | 0.65 |
| ML735004 | 154316 | 156371 | BDV34DRAFT_228389 | major facilitator superfamily domain-containing protein | 0 | 0.9 | 0.81 |
| ML735005 | 75002 | 75586 | BDV34DRAFT_228432 | Carboxypeptidase Y | 0 | 0.8 | 0.65 |
| ML735007 | 22963 | 23411 | BDV34DRAFT_215627 | DUF953 domain-containing protein | 0.1 | 1 | 0.81 |
| ML735007 | 90578 | 95624 | BDV34DRAFT_157380 | TPR_REGION domain-containing protein | 0 | 0.8 | 0.65 |
| ML735009 | 102819 | 105071 | BDV34DRAFT_201533 | cytochrome P450 monooxygenase | 0 | 0.7 | 0.51 |
| ML735012 | 15639 | 16747 | BDV34DRAFT_237217 | CBAH domain-containing protein | 0 | 0.9 | 0.81 |
| ML735012 | 18081 | 18703 | BDV34DRAFT_201837 | HNH endonuclease | 0 | 0.9 | 0.81 |
| ML735012 | 18979 | 20709 | BDV34DRAFT_201838 | Solute carrier family 40 protein | 0 | 0.9 | 0.81 |
| ML735012 | 21670 | 24108 | BDV34DRAFT_228850 | Protein kinase domain-containing protein | 0 | 1 | 1.00 |
| ML735012 | 24235 | 26453 | BDV34DRAFT_237219 | FAD-binding PCMH-type domain-containing protein | 0 | 1 | 1.00 |
| ML735012 | 26680 | 30135 | BDV34DRAFT_237221 | Fungal_trans domain-containing protein | 0 | 1 | 1.00 |
| ML735012 | 31401 | 31976 | BDV34DRAFT_228854 | Phage portal protein | 0 | 1 | 1.00 |
| ML735012 | 32064 | 32537 | BDV34DRAFT_228855 | AMP-binding domain-containing protein | 0 | 1 | 1.00 |
| ML735012 | 33304 | 33726 | BDV34DRAFT_228856 | DUF418 domain-containing protein | 0 | 1 | 1.00 |
| ML735012 | 34598 | 36532 | BDV34DRAFT_237223 | major facilitator superfamily domain-containing protein | 0 | 1 | 1.00 |
| ML735012 | 36910 | 37284 | BDV34DRAFT_228858 | Transposase | 0 | 1 | 1.00 |
| ML735012 | 37934 | 39498 | BDV34DRAFT_237226 | RmlC-like cupin domain-containing protein | 0 | 1 | 1.00 |
| ML735012 | 40901 | 42143 | BDV34DRAFT_215917 | GH131_N domain-containing protein | 0 | 1 | 1.00 |
| ML735014 | 34248 | 34560 | BDV34DRAFT_228962 | Aminotran_1_2 domain-containing protein | 0 | 0.8 | 0.65 |
| ML735014 | 34907 | 35855 | BDV34DRAFT_228963 | PAS domain-containing protein | 0 | 0.9 | 0.81 |
| ML735016 | 50708 | 53032 | BDV34DRAFT_216106 | nucleoside phosphorylase domain-containing protein | 1 | 0.1 | 0.81 |
| ML735019 | 117613 | 118773 | BDV34DRAFT_216281 | F-box domain-containing protein | 0 | 0.7 | 0.51 |
| ML735019 | 119188 | 119627 | BDV34DRAFT_229268 | Bacteriophage protein | 0 | 0.7 | 0.51 |
| ML735022 | 51385 | 53456 | BDV34DRAFT_202966 | Zn(2)-C6 fungal-type domain-containing protein | 0 | 0.8 | 0.65 |
| ML735022 | 53677 | 55406 | BDV34DRAFT_202967 | aromatic compound dioxygenase | 0 | 0.8 | 0.65 |
| ML735024 | 94661 | 97299 | BDV34DRAFT_229521 | fungal-specific transcription factor domain-containing protein | 0.3 | 1 | 0.51 |
| ML735024 | 97793 | 98936 | BDV34DRAFT_216497 | kinase domain protein | 0 | 1 | 1.00 |
| ML735025 | 53881 | 54385 | BDV34DRAFT_229547 | unspecified product | 0 | 0.9 | 0.81 |
| ML735025 | 90352 | 91957 | BDV34DRAFT_229554 | MFS general substrate transporter | 0.1 | 0.9 | 0.62 |
| ML735027 | 12884 | 13675 | BDV34DRAFT_229613 | DUF4294 domain-containing protein | 0 | 1 | 1.00 |
| ML735027 | 15623 | 18988 | BDV34DRAFT_216570 | glycosyl hydrolase family 71-domain-containing protein | 0 | 1 | 1.00 |
| ML735028 | 27804 | 29649 | BDV34DRAFT_179322 | Allantoinase | 2 | 1 | 1.00 |
| ML735033 | 970 | 1338 | BDV34DRAFT_229870 | WG repeat-containing protein | 0 | 0.9 | 0.81 |
| ML735033 | 1501 | 3348 | BDV34DRAFT_229871 | tubby C-terminal-like domain-containing protein | 0 | 0.9 | 0.81 |
| ML735033 | 3411 | 5383 | BDV34DRAFT_216794 | aspartic peptidase domain-containing protein | 0 | 1 | 0.70 |
| ML735033 | 9474 | 11561 | BDV34DRAFT_182674 | Transferred entry: 2.1.1.354 | 0 | 0.8 | 0.65 |
| ML735033 | 11775 | 12560 | BDV34DRAFT_229875 | P-loop containing nucleoside triphosphate hydrolase protein | 0 | 0.8 | 0.65 |
| ML735033 | 89090 | 92026 | BDV34DRAFT_237931 | acetyl-CoA synthetase-like protein | 0 | 0.8 | 0.65 |
| ML735033 | 94353 | 96196 | BDV34DRAFT_216823 | cytochrome P450 | 0 | 0.8 | 0.65 |
| ML735033 | 97231 | 98807 | BDV34DRAFT_229908 | terpenoid synthase | 0 | 0.8 | 0.65 |
| ML735033 | 99385 | 101220 | BDV34DRAFT_237932 | cytochrome P450 monooxygenase | 0 | 0.8 | 0.65 |
| ML735033 | 102514 | 103543 | BDV34DRAFT_216826 | Cyclopropane-fatty-acyl-phospholipid synthase | 0 | 0.8 | 0.65 |
| ML735035 | 8773 | 9470 | BDV34DRAFT_204045 | Secreted protein | 0 | 0.7 | 0.51 |
| ML735035 | 53579 | 54655 | BDV34DRAFT_204069 | EamA domain-containing protein | 0 | 0.8 | 0.65 |
| ML735035 | 56361 | 57616 | BDV34DRAFT_232043 | ATP-grasp domain-containing protein | 0 | 0.8 | 0.65 |
| ML735035 | 58368 | 59908 | BDV34DRAFT_232045 | Kama family protein | 0 | 0.8 | 0.65 |
| ML735035 | 60967 | 63480 | BDV34DRAFT_216885 | Zn(2)-C6 fungal-type domain-containing protein | 1 | 0.3 | 0.51 |
| ML735035 | 67502 | 67825 | BDV34DRAFT_229977 | UDENN domain-containing protein | 0 | 1.2 | 0.65 |
| ML735046 | 97673 | 100020 | BDV34DRAFT_205037 | homeobox KN domain-containing protein | 1.9 | 1 | 0.81 |
| ML735046 | 100760 | 101203 | BDV34DRAFT_205038 | unspecified product | 1.9 | 1 | 0.81 |
| ML735050 | 73288 | 74994 | BDV34DRAFT_232415 | PA14 domain-containing protein | 0 | 0.9 | 0.81 |
| ML735050 | 76374 | 77094 | BDV34DRAFT_205351 | Ovule protein | 0 | 0.8 | 0.65 |
| ML735050 | 87083 | 88201 | BDV34DRAFT_230515 | Solute carrier family 40 protein | 0 | 0.8 | 0.65 |
| ML735075 | 23341 | 24078 | BDV34DRAFT_231190 | Rho-GAP domain-containing protein | 0 | 0.9 | 0.81 |
| ML735075 | 24314 | 26196 | BDV34DRAFT_39909 | major facilitator superfamily domain-containing protein | 0 | 0.9 | 0.81 |
| ML735075 | 26441 | 28830 | BDV34DRAFT_231192 | C2H2 type zinc finger domain protein | 0 | 0.9 | 0.81 |
| ML735075 | 28994 | 30537 | BDV34DRAFT_233053 | cytochrome P450 | 0 | 0.9 | 0.81 |
| ML735075 | 39131 | 40267 | BDV34DRAFT_218026 | Zn(2)-C6 fungal-type domain-containing protein | 0 | 0.9 | 0.81 |
| ML735075 | 44094 | 46419 | BDV34DRAFT_231196 | cytochrome P450 | 0 | 1.8 | 0.61 |
| ML735075 | 46633 | 49517 | BDV34DRAFT_233055 | Zn(2)-C6 fungal-type domain-containing protein | 0 | 1.5 | 0.81 |
| ML735075 | 49882 | 51054 | BDV34DRAFT_233057 | RmlC-like cupin domain-containing protein | 0 | 1.5 | 0.81 |
| ML735075 | 51313 | 52664 | BDV34DRAFT_231199 | LEA_2 domain-containing protein | 0 | 1 | 1.00 |
| ML735075 | 54611 | 55693 | BDV34DRAFT_231200 | Amidohydro-rel domain-containing protein | 0.2 | 1 | 0.65 |
| ML735085 | 42217 | 42789 | BDV34DRAFT_207245 | Transcriptional regulator | 0 | 1 | 1.00 |
| ML735093 | 16907 | 20215 | BDV34DRAFT_218307 | RING-type domain-containing protein | 0 | 0.7 | 0.51 |
| ML735093 | 22979 | 23829 | BDV34DRAFT_231500 | tryptophan dimethylallyltransferase-domain-containing protein | 0 | 0.7 | 0.51 |
| ML735094 | 32064 | 32786 | BDV34DRAFT_231515 | Protein Ves | 0 | 0.7 | 0.51 |
| ML735119 | 10645 | 14158 | BDV34DRAFT_231724 | MACPF domain-containing protein | 0 | 0.7 | 0.51 |

*Scaff=scaffold ID, gene_start=coordinate of start site, gene_stop=coordinate of stop site, gene_ID=gene ID relative to *A. parasiticus* CBS-117618, gene anno=functional prediction via UniProt, Av AS=average CN in AS, Av AP=average CN in AP, V_ST_=V_ST_ value
